# Copy number variants alter local and global mutational tolerance

**DOI:** 10.1101/2022.12.30.521611

**Authors:** Grace Avecilla, Pieter Spealman, Julia Matthews, Elodie Caudal, Joseph Schacherer, David Gresham

## Abstract

Copy number variants (CNVs), duplications and deletions of genomic content, contribute to evolutionary adaptation, but can also confer deleterious effects, and cause disease. Whereas the effects of amplifying individual genes or whole chromosomes (i.e., aneuploidy) have been studied extensively, much less is known about the genetic and functional effects of CNVs of differing sizes and structures. Here, we investigated *Saccharomyces cerevisiae* (yeast) strains that have CNVs of variable structures but with multiple copies of the gene *GAP1*. Although beneficial in glutamine-limited chemostats, CNVs result in decreased fitness compared with the euploid ancestor in rich media. We used transposon mutagenesis to investigate mutational tolerance and genetic interactions with CNVs. We find that CNVs confer novel mutational tolerance in amplified essential genes and novel genetic interactions. We validated a novel genetic interaction with *BMH1*. CNV strains have increased mutational tolerance in genes related to translation, and reduced mutational tolerance in genes related to mitochondrial function. We performed RNAseq and found that transcriptional dosage compensation does not affect the majority of genes amplified by CNVs. Furthermore, we do not find that CNV strains exhibit previously described transcriptional signatures of aneuploidy. Instead, CNV strains exhibit downregulation of genes involved in cellular respiration, nucleoside biosynthetic processes, and small molecule metabolism, and upregulation of genes involved in transposition, nucleic acid metabolic processes, and siderophore transport. Our study reveals the extent to which local and global mutational tolerance is modified by CNVs with implications for genome evolution and CNV associated diseases, such as cancer.

## Introduction

Evolution occurs through changes to an organism’s genome and selection on the functional effects of those changes. Genomes can evolve in many ways, including through single nucleotide changes, structural rearrangements, deletion and duplication of segments of DNA. Amplification of segments of DNA, a type of copy number variation (CNV), is a major force in rapid adaptive evolution and genome evolution. In the short term, gene amplification can result in increased gene expression which provides a selective advantage facilitating rapid adaptive evolution (Kondrashov 2012; Myhre et al. 2013). In the long term, amplification of genes may relax selective constraints, allowing accumulation of mutations on the additional gene copies and gene evolution through subfunctionalization or neofunctionalization (Ohno 2014; Freeling et al. 2015; Innan and Kondrashov 2010). Rapid adaptation through gene amplification is prevalent throughout the tree of life. In particular, gene amplification has been shown to mediate rapid adaptation to a variety of selective pressures from nutrient limitation to antibiotics in both natural and experimental populations of microbes (Lauer et al. 2018; Selmecki et al. 2009; Todd and Selmecki 2020; Gresham et al. 2008; Hong and Gresham 2014; Dhami et al. 2016; Nair et al. 2008; Pränting and Andersson 2011; Paulander et al. 2010). Gene amplification is also common in cancers, and can promote tumorigenesis (Ben-David and Amon 2020). Oncogene amplification confers enhanced proliferation properties to cells driving their aberrant growth. Thus, understanding the evolutionary, genetic, and functional consequences of CNVs is of central importance. The budding yeast, *Saccharomyces cerevisiae*, has been extensively used as a model to study the effects of gene amplification on cellular properties, fitness, and genetics.

Copy number variants can range from small duplications and deletions to the gain or loss of whole chromosomes, known as aneuploidy. Previous studies have investigated the effect of amplifying individual genes primarily using plasmid libraries with native or inducible promoters (Moriya 2015). These studies have found that in commonly used laboratory strains around 10-20% of genes are deleterious whereas 0-5% are beneficial when overexpressed (Ascencio et al. 2021; Sopko et al. 2006; Arita et al. 2021; Douglas et al. 2012). These effects are dependent on genetic background as a recent study found significant variation in the number of genes that are deleterious when overexpressed in 15 genetically diverse yeast lineages (Robinson et al. 2021). Fitness effects of gene amplification tend to be dependent on both the particular gene amplified and the environmental context, though most amplified genes have neutral effects regardless of environment (Ascencio et al. 2021; Payen et al. 2016). To date, there is conflicting evidence for the Dosage Balance hypothesis, which predicts that genes whose products are involved in protein complexes or have many interactions are more likely to be deleterious when overexpressed due to stoichiometric imbalances (Birchler and Veitia 2012; Rice and McLysaght 2017). Some studies find that genes that are deleterious when overexpressed are enriched for protein complexes and protein interactions (Robinson et al. 2021; Makanae et al. 2013), whereas others studies do not (Ascencio et al. 2021; Sopko et al. 2006; Arita et al. 2021). Single gene overexpression libraries have also been used to identify synthetic dosage lethal interactions with gene deletions (Douglas et al. 2012; Sopko et al. 2006; Liu et al. 2009) in which an overexpressed gene is deleterious in the background of a gene knock out.

Aneuploidy is frequent (appr. 20%) in strains of yeast isolated from diverse ecologies (Peter et al. 2018; Hose et al. 2015; Gallone et al. 2016; Zhu et al. 2016; Scopel et al. 2021), and these aneuploids grow similarly to their euploid counterparts (Hose et al. 2015; Gasch et al. 2016). Aneuploids also frequently arise in evolution experiments and are associated with increased fitness (Sunshine et al. 2015; Lauer et al. 2018; Hong and Gresham 2014; Rancati et al. 2008; Gresham et al. 2008; Yona et al. 2012). However, adaptive aneuploids can exhibit antagonistic pleiotropy such that they are deleterious in other environments (Sunshine et al. 2015; Linder et al. 2017). Seminal studies of one laboratory strain, W303, found that aneuploids grow more slowly than euploids, regardless of karyotype (Torres et al. 2007; Sheltzer et al. 2012; Beach et al. 2017), exhibit a transcriptional signature characteristic of the yeast environmental stress response (ESR) (Torres et al. 2007; Sheltzer et al. 2012), and result in a variety of cellular stresses, including proteotoxic, metabolic, and mitotic stress (reviewed in (Zhu et al. 2018)). These effects may also be background dependent as a recent study mapped differences in aneuploidy tolerance between W303 and wild yeast strains to a single gene, *SSD1*, which has a truncating mutation in W303 (Hose et al. 2020). *SSD1* is a RNA-binding translational regulator, whose targets include mitochondrial transcripts. Loss of *SSD1* function results in defects in mitochondrial function and proteostasis that enhance sensitivity to aneuploidy (Hose et al. 2020). In addition to observing different fitness effects, studies of aneuploids in different genetic backgrounds have found differing results in transcriptomic dosage compensation, ESR, and proteotoxic stress (Muenzner et al.; Torres et al. 2007; Pavelka et al. 2010; Larrimore et al. 2020; Dephoure et al. 2014; Hose et al. 2015; Gasch et al. 2016; Zhu et al. 2018).

Whereas numerous studies have investigated the effects of single gene amplifications and whole chromosome aneuploidy, little is known about the effects of CNVs that vary in size and structure. One survey sought to study the fitness effects of a diverse set of synthetic amplicons extending from the telomere and ranging in size from 0.4 to 1,000 kb across the genome in diploid yeast and measured their fitness in three conditions (Sunshine et al. 2015). Through comparison to single-gene amplifications (Payen et al. 2016), they found that the distribution of fitness effects for telomeric amplicons was broader than that of single gene amplifications. Notably, they also found that of the telomere amplified regions that affected fitness, 94% had condition-dependent effects. However, it remains unknown whether there are common fitness effects, genetic interactions, or transcriptomic states associated with non-engineered CNVs.

In this study, we investigated seven yeast strains containing diverse CNV structures. The strains all contain amplification of the gene *GAP1* and were previously isolated from evolution experiments in glutamine-limited chemostats (Lauer et al. 2018). We found that despite having fitness greater than or equal to the ancestral euploid in glutamine-limited chemostats, most CNV-containing lineages have fitness defects in rich media with galactose as a carbon source. We used transposon mutagenesis to investigate how CNVs alter mutational tolerance within the copy number altered region and throughout the rest of the genome. We find that CNVs relieve gene essentiality enabling the acquisition of genetic variation in genes that are essential when present as a single copy. We find evidence for extensive extragenic genetic interactions with CNVs that are both common and strain specific. We investigated how CNVs alter the transcriptome, and found that while amplified genes do have increased mRNA expression, some strains appear to exhibit dosage compensation. We did not observe previously described transcriptional signatures of aneuploidy in CNV strains. Instead, we find that CNV-containing strains tend to have decreased expression of genes involved in respiration, nucleoside biosynthetic processes, and small molecule metabolism, and increased expression of genes involved in transposition, nucleic acid metabolic processes, and siderophore transport. Taken together, our experiments suggest there are both common and strain specific interactions and transcriptional responses that affect fitness in yeast with CNVs.

## Results

### *GAP1* CNVs confer variable fitness effects

Previously, we performed experimental evolution using budding yeast cells in glutamine-limited chemostats for approximately 270 generations (Lauer et al. 2018). The yeast strain (a haploid derivative of S288C) used to inoculate the evolution experiments contained a fluorescent reporter gene adjacent to the general amino acid permease gene, *GAP1*, on chromosome XI. The proximity of the fluorescent gene (~1 kilobase from the 5’ end of *GAP1*) ensures that it is typically co-amplified with *GAP1* enabling efficient detection and isolation of CNV-containing strains. The structures of some *GAP1* CNVs are complex and can only be imperfectly estimated using short-read sequencing and pulse-field electrophoresis (Lauer et al. 2018). Therefore, we resolved the structures of the seven CNV strains using long-read sequencing and genome assembly (Spealman et al. 2022) to enable accurate estimation of gene copy number within the CNV. The resolved CNV structures (**Figure 1A; SFig 1; STable 1**) range in *GAP1* copy number from two (Aneu), to three (Trip1, Trip2, ComTrip, Sup), to four (ComQuad, ComSup) copies; in the number of amplified genes from 20 to 315; and in the total amount of amplified DNA from ~79,000 to ~667,000 additional nucleotides (**STable 2; STable 3**). The *GAP1* CNVs have a variety of structures, including an aneuploid (Aneu), inverse triplications characteristic of origin dependent inverted repeat amplification (ODIRA; Trip1, Trip2), a supernumerary chromosome (Sup) and more complex structures probably resulting from multiple mutational events (ComTrip, ComQuad, ComSup). In addition to *GAP1* CNVs, each strain contains a small number of unique nucleotide variants compared to the ancestor and two strains also contain independant amplifications at the *GLC7* locus on chromosome V (**STable 4; SFig 1**).

**Figure 1.**
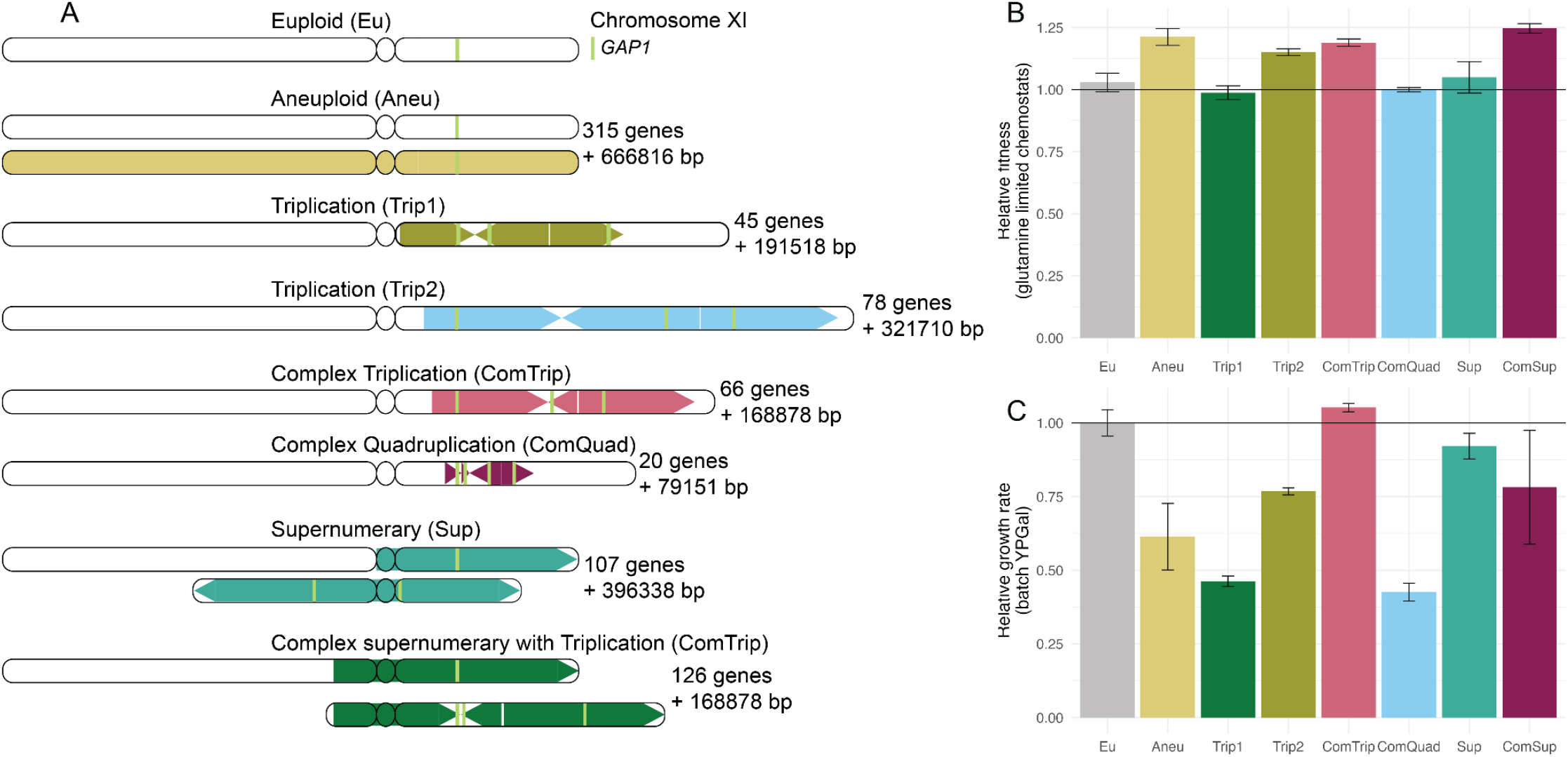
Strains with *GAP1* CNVs differ in structure and fitness. **A)** We previously evolved a euploid *S. cerevisiae* strain in glutamine-limited chemostats and isolated seven strains that have CNVs on Chromosome XI that include *GAP1 (Lauer et al. 2018*). The structure of each *GAP1* CNV was resolved using long read sequencing (Spealman et al. 2022) and is summarized here. The amplified region is shown as a colored block with arrows. Arrows pointing right represent copies that maintain their original orientation, whereas arrows pointing left represent copies that are inverted. The number of genes amplified and the number of additional base pairs (bp) are annotated. **B)** The relative fitness (compared to the ancestral strain) of evolved strains containing *GAP1* CNVs was determined by pairwise competition experiments with a nonfluorescent reference strain in glutamine-limited chemostats. Error bars are 95% confidence intervals for the slope of the linear regression. **C)** Average and standard deviation (error bars) growth rate relative to the ancestral, euploid strain in YPGal batch culture. Horizontal black lines in **B** and **C** denote the ancestral euploid fitness.

All CNV strains have fitness greater than or equal to the ancestral euploid strain in the glutamine-limited environment in which they evolved (**Figure 1B**). However, the CNV strains grow slower than the euploid strain in a non-selective environment: yeast-peptone-galactose (YPGal) batch culture (**Figure 1C**). The fitness benefit in the environment in which they evolved and the fitness deficit in the alternative environment differ between strains. Fitness benefits and costs do not correlate with the number of additional bases or the number of open reading frames in the CNV region (**SFig 2**).

### Transposon mutagenesis reveals tolerance to mutation

We sought to investigate the genetic impact of CNVs using high-throughput genetics. Previous studies using transposon mutagenesis in bacteria and yeast have shown that transposon insertion density reflects tolerance to mutation and is an efficient means of identifying genomic regions essential for cell survival in a specific environment or genetic background (Michel et al. 2017; Guo et al. 2013; Grech et al. 2019; Segal et al. 2018; Gale et al. 2020; Levitan et al. 2020). We generated *Hermes* insertion libraries in each CNV strain and in two independent replicates of the ancestral euploid strain using modifications of published methods (Gangadharan et al. 2010; Caudal et al.) (**Figure 2A**; **Materials and Methods**). Briefly, *Hermes* transposition was induced in YPGal media using batch cultures undergoing serial transfer, and transposition events were selected using an antibiotic marker. Insertion sites were identified by targeted PCR, followed by library preparation and deep sequencing (**Materials and Methods**). Unique insertion sites and the number of reads per insertion site were identified using a custom bioinformatic pipeline (**Materials and Methods, STable 5**). As our sequencing and analysis pipeline cannot differentiate between identical sequence reads that result from unique priming events from those due to PCR duplicates, we quantified the number of unique insertion sites per gene, unless otherwise noted (**STable 6; STable 7)**. The nine libraries exhibited variation in the number of unique insertion sites which scaled with the total number of reads sequenced (**SFig 3; STable 8, STable 9**). To normalize for differences in sequencing depth, we determined the number of insertions per million reads for analyses (**Materials and Methods**).

**Figure 2.**
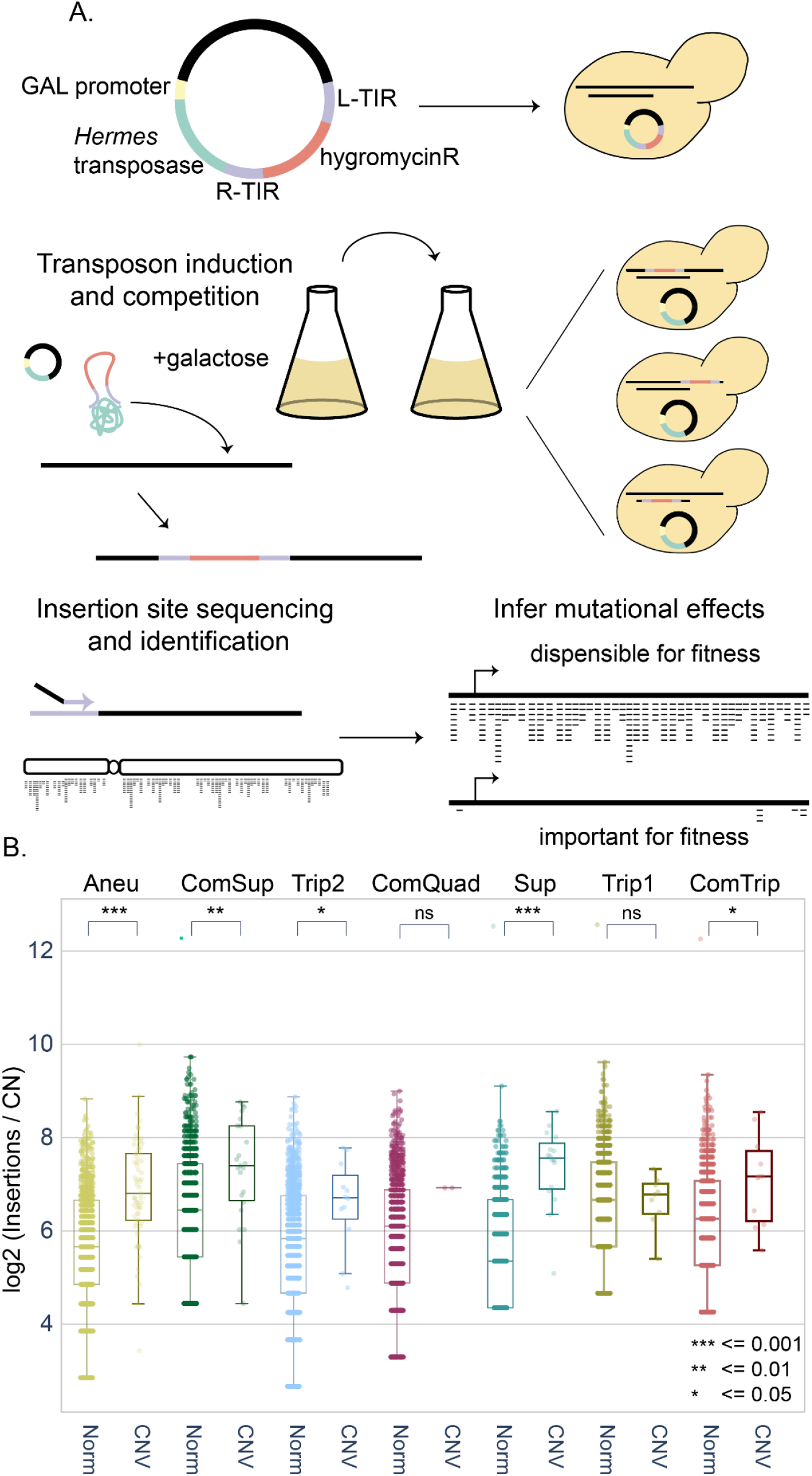
Profiling mutation tolerance in CNV strains using insertional mutagenesis. **A)** Plasmids containing the hermes transposase regulated by the GALS promoter (a truncated GAL1 promoter) and a hygromycin resistance gene flanked by the hermes terminal inverted repeats (TIR) were transformed into each yeast strain. Upon addition of galactose, the transposase is expressed, and the hygromycin resistance gene flanked by the TIRs is excised from the plasmid and inserted in the yeast genome. DNA is extracted, digested with restriction enzymes, and circularized. Insertion sites are identified by inverse PCR and amplicon sequencing. Mutational tolerance is inferred by the number of unique insertion sites at a given region of the genome. **B)** Unique insertion sites per gene copy number for essential genes (Winzeler et al. 1999). Genes are defined as either CNV associated (CNV) or not (Norm). A Mann-Whitney U statistic was used to compare distributions between the CNV and Norm essential genes, P-values are indicated by the following: ns: p > 0.05; * <= 0.05, ** <= 0.01, *** <= 0.001

As our transposition protocol entails serial propagation with periodic bottlenecking, the mutation frequency per gene reflects the tolerance to mutation in the rich media condition. We compared our transposon insertion data to a list of essential genes generated in YPD (Winzeler et al. 1999), and to relative fitness measurements of genes grown on YPGal (Costanzo et al. 2021), that were defined using complete open reading frame deletions. In all CNV and euploid strains, essential genes have fewer insertions than non-essential genes (**SFig 4, SFig 5**). GAL genes in the CNV strains also had no significant change in insertions relative to the euploid. These results confirm that transposon insertion density is a reliable predictor of sequence tolerance to disruptive mutation in CNV strains.

### Gene amplification increases mutational target size

We investigated how gene amplification affects insertion density by considering only coding sequences (which we refer to as genes) within the CNV region for each CNV strain, and comparing insertion density with all genes on chromosome XI in the euploid replicates. We find that in 6 of 7 CNV strains, amplified genes have a higher insertion frequency than in the euploid (paired t-test, p<0.0001), consistent with increased target size resulting in increased mutation frequency. The single exception, ComQuad, is likely due to insufficient statistical power, since it has the fewest amplified genes.

Essential genes (Winzeler et al. 1999) have significantly fewer insertions than non-essential genes in the euploid strain and in unamplified genes in the CNV strains (**SFig 8A,** Welch’s two sample t-test, p<0.0001). To account for the differences in copy number, we normalized insertion frequency by gene copy number. Amplified essential genes within CNVs have significantly higher (Mann-Whitney U, p<=0.05) copy-number corrected insertion frequencies than essential genes in unamplified regions of the genome for 5 of 7 CNV strains consistent with relaxed selection for essentially upon amplification. The two exceptions, ComQuad and Trip1, have the fewest essential genes within the CNV and thus are likely underpowered for observing this effect. Mutation frequencies in essential genes within the CNV are also increased in these strains. (**Figure 2B**).

To study the impact of CNVs on mutational tolerance across the entire genome, we compared genome-wide mutation frequencies in the CNV strains with the euploid ancestor. The number of insertions in a gene in the euploid strains is positively correlated with the number of insertions in a CNV strain (**SFig 6**), (**SFig 7)**. The strength of the relationship, assessed using the slope of the regression line, is less than that expected on the basis of copy number for all strains. However, this is likely due to a compositional data effect, as the slope of regression performed on unamplified genes in most CNV strains is slightly less than one.

The number of insertions in some amplified genes is not well predicted by the model (i.e., they have large residuals); when the number of insertions is higher than the predicted value, this may indicate a gene that is particularly sensitive to amplification. We also found a significant enrichment in outliers (genes with standardized residuals > 2) in the CNV associated genes relative to the background (FET, p <= 0.05), consistent with amplified genes having more insertions.

For example, *UTH1* is amplified in all CNV strains and is a significant outlier in each (S**Table 10, Stable 11)** suggesting that increased *UTH1* copy number is deleterious. *UTH1*, has been shown to lead to cell death when overexpressed in the W303 genetic background (Camougrand et al. 2003). *UTH1* is a mitochondrial protein involved in regulating both mitochondrial biogenesis and degradation (Camougrand et al. 2004), and is regulated by *SSD1*, whose loss of function is associated with fitness defects in aneuploid strains of W303 (Hose et al. 2020). Two other genes, *VPS1* and *ECM9*, also exhibit this same pattern although they are only amplified in 3 and 4 strains, respectively. Each of these is enriched in hits, suggesting they are DSGs. Only one non-CNV gene has a similar profile, *PDR5* which is significantly enriched in 4 of the 7 strains.

Conversely, significantly reduced insertional frequency may reflect an advantage due to increased copy number. For example, *YKR005C* is significantly depleted in expected mutation frequency in three of four CNV strains in which it is amplified (**STable 10)**.

We also observe genes throughout the genome that have significantly higher rates of insertion in the euploid ancestor than CNV strains. For example, *GPB1* (YOR371C) and *GPB2* (YAL056W) have significantly higher mutation frequencies in the ancestor relative to every CNV strain (S**Table 10)**. These paralogs are multistep regulators involved in the cAMP-PKA signaling and RAS signaling pathways. Interestingly, the 18 genes with significantly higher mutation frequencies in the ancestor relative to the majority of the evolved strains are enriched for functions in GTPase activity (GO:0007264), negative regulation of RAS (GO:0046580), and negative regulation of cAMP (GO:2000480), (GSEA, adj.p-value<=0.05). This suggests that the RAS/PKA pathway may enhance tolerance of CNVs.

### CNVs result in common and strain specific genetic interactions

Genes that have no insertions events may be essential and intolerant of mutation, or may have no insertions due to chance. To establish a genome-wide view of differences in mutational tolerance between CNV strains and the euploid strain, we first identified 327 genes that have no insertions in either replicate of the euploid strain. Of these, 136 (42%) have previously been annotated as essential or as having low fitness in galactose (S**Table 12**). We define the set of 327 genes as “euploid intolerant”. Many of these genes had insertions in one or more of the CNV strains and seven euploid intolerant genes had insertions in all CNV strains (**Figure 3A**). These seven genes have all been previously annotated as essential or as having low fitness in galactose. Although four of these genes were amplified in one or more CNV strains, none are amplified in all CNV strains (**Figure 3A**), suggesting that mutational tolerance in the CNV strains is not simply attributable to increased target size. We also looked for genes that had no insertions in any CNV strain but did have insertions in both replicates of the euploid strain: we identified one gene, *RRN10*. However, it only had one insertion in Eu replicate1 and two insertions in Eu replicate 2, so this is unlikely to be meaningful. This suggests that CNVs do not confer novel genetic vulnerabilities.

**Figure 3.**
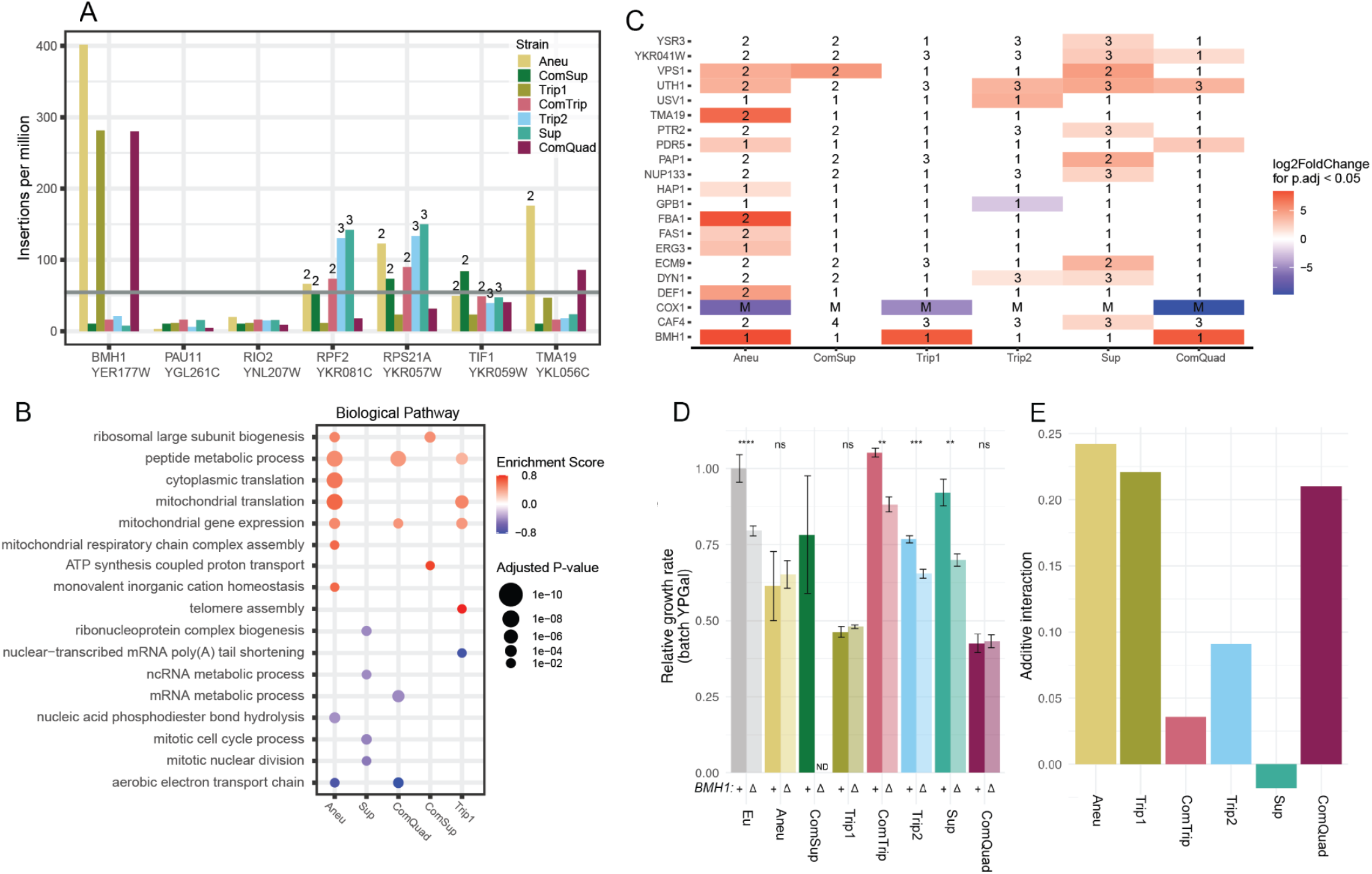
CNV strains have common and allele specific genetic interactions. **A)** Seven genes have no insertions in either replicate of the euploid strain, whereas insertions are identified in these genes in all CNV strains. The gray line represents the median insertions per million per gene across all strains. Numbers indicate the copy number if a gene is contained within the CNV. **B)** Enriched GO terms identified using Gene Set Enrichment Analysis (GSEA). GSEA was applied to a ranked gene list based on log2 fold changes obtained in differential analysis comparing each CNV insertion profile to the euploid insertion profiles, with the false discovery rate (FDR) for enriched terms set to 0.05. Terms with adjusted p-value < 0.05 are shown (circle size). Positive enrichment scores (red) indicate functions that have increased insertions in the CNV strain. Negative enrichment scores (blue) indicate functions that have decreased insertion frequencies in the CNV strain. Neither Trip2 or ComTrip had significant enrichments of any gene sets **C)** Significant genes (p.adjust<0.05) from differential analysis comparing each CNV insertion profile to the euploid insertion profiles. Positive log2 Fold Change values have more insertions in CNV strains than euploid strains, whereas negative log2 Fold Change values indicate genes with fewer insertions in the CNV strain than euploid strains. If a gene is amplified the copy number is annotated. **D)** Average and standard deviation (error bars) of growth rate relative to the ancestral, euploid strain in YPGal batch culture. P-values from two-sample t-test are indicated by the following: ns: not significant; *: p < 0.05; **: p < 0.01; ***: p < 0.001; ****: p < 0.0001. **E)** Additive genetic interaction (epsilon = double - single*single) for CNV and *BMH1* double mutants, calculated from growth rates shown in (**D**).

To quantitatively assess genetic interactions between the CNVs and all other genes throughout the genome, we quantified differential insertion frequency using the number of unique insertion sites per gene between each CNV strain and the euploid replicates (S**Fig 6**). To assess the global trend, we performed gene set enrichment analysis (GSEA) using the ranked list of fold change in number of insertions (**Figure 3B; STable 13**). We found that three strains, Aneu, ComQuad, and Trip1, have an increased mutational tolerance in genes annotated with terms related to translation and mitochondrial gene expression, and decreased mutational tolerance for genes with functions in the aerobic electron transport chain. We also observed enrichment for terms that are unique to individual strains. For example, the Supernumerary chromosome (Sup) exhibits decreased tolerance for mutations in genes with functions in the mitotic cell cycle and nuclear division (**Figure 3B**).

We identified individual genes with differences in insertional tolerance in CNV strains compared to the ancestral euploid strain (**Figure 3C, STable 14**). Most genes that are significant in one strain tend to have similar trends in other CNV strains, with few exceptions (S**Fig 9A**). The Aneu, ComQuad, and Trip1 strains all have significantly more insertions than the euploid ancestor in *BMH1*, which is involved in many processes including regulation of mitochondrial-nuclear signaling (Liu et al. 2003), carbon metabolism (Dombek et al. 2004), as well as transcription and chromatin organization (Kumar 2017; Jain et al. 2021). These strains also have significantly fewer insertions in genes encoded by the mitochondrial genome, including *COX1*. Interestingly, we found these strains also showed greatly reduced growth when plated on glycerol media, suggesting impaired mitochondrial function or disregulation in retrograde signaling (Roca-Portoles and Tait 2021).

Differences in insertion tolerance in genes that are not contained within the CNV reflect differential genetic interactions. To confirm this, we generated complete deletions of the coding sequence of *BMH1* in all strains except ComSup, for which we could not obtain a transformant, and measured growth rates of the single and double mutants in YPGal (**Figure 3D**). We find that deletion of *BMH1* does not significantly affect growth rate in Aneu, ComQuad, and Trip1, but does result in reduced growth rate in other strains. We calculated the genetic interaction of *BMH1* with the CNV for each strain (Mani et al. 2008), and confirmed positive interactions for these three strains, consistent with transposon insertion profiles (**Figure 3E**, **SFig 9B**).

### Amplified genes have increased mRNA expression

To test how gene expression impacts mutational tolerance in CNV lineages, we performed RNAseq in triplicate on each euploid and CNV strain growing in YPGal, and quantified gene expression in each CNV strain relative to the euploid ancestor (S**Table 15**, **STable 16, STable 17**). First, we investigated genes encoded on chromosome XI for evidence of dosage compensation within the CNV region (**STable 18)**. We found that in each CNV strain, amplified genes have significantly higher mRNA expression than in the euploid ancestor (Mann-Whitney U p < 0.001, **Figure 4A, STable 19**), and expression in amplified genes is highly correlated with euploid expression (S**Fig 10**) with many amplified genes being significantly higher than their euploid counterparts (**STable 18**). After correcting for copy-number (**STable 20**), we find the RNA expression is in agreement with the abundances observed in the ancestor (**Figure 4B, STable 21, STable 22**), suggesting that dosage compensation does not operate in these strains under these conditions.

**Figure 4.**
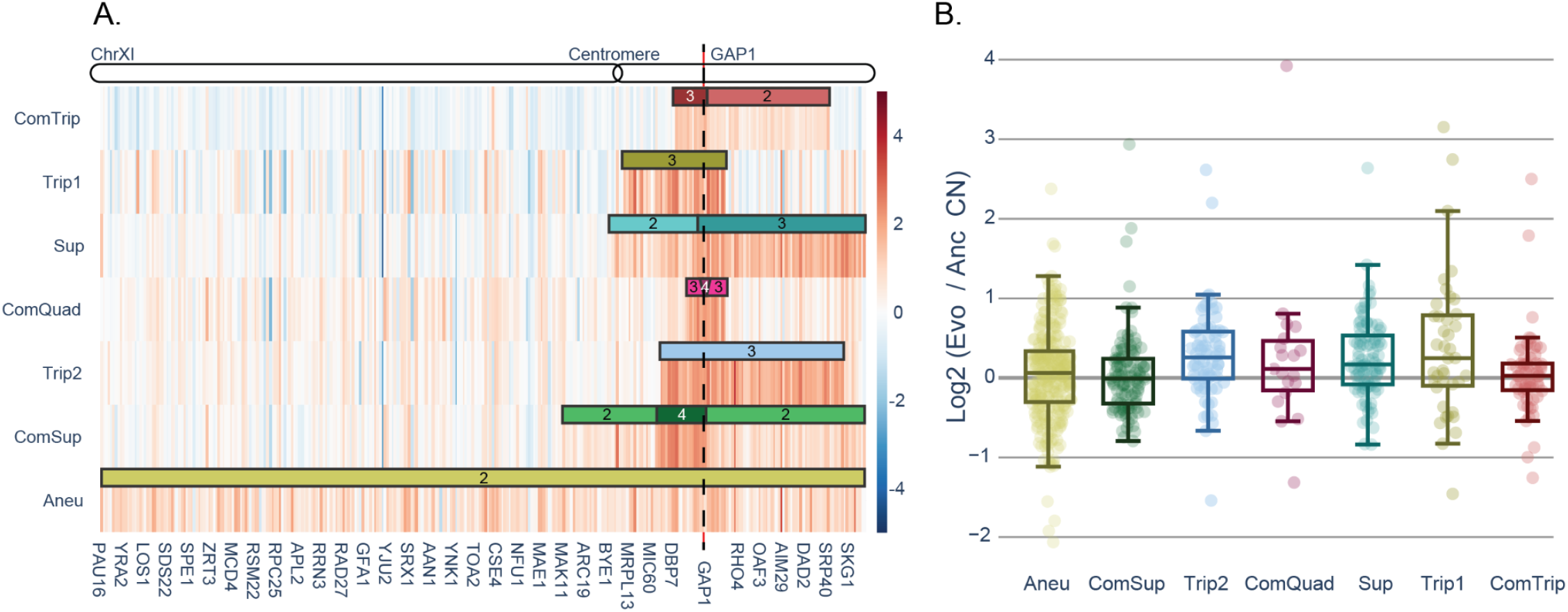
Amplified genes result in increased mRNA expression but are not associated with increased mutation frequency. **A)** Replicate averaged log2 fold change of all genes on chromosome XI, ordered as on the chromosome, in each CNV strain compared to the euploid ancestor. Black lines denote CNV boundaries. Numbers indicate the copy number of genes within the CNV. **B)** log2 fold change mRNA expression compared to the euploid for genes that are amplified in each CNV strain normalized by gene copy number..

To test the relationship between gene expression and mutational tolerance, we compared the log2 fold change of transposon insertion frequency with the log2 fold change for mRNA expression for each CNV strain compared to the euploid strain. If CNV burden is related to a general cost associated with increased expression of all amplified genes, or the “mass action” of the CNV, we expect that the fold change in transposon insertions would positively correlate with the fold change in mRNA expression. However, we do not observe a significant correlation between mRNA abundance and transposon insertion in any of the strains (p > 0.05) (SFig 11). This supports the hypothesis that adverse effects of CNVs do not stem from the “mass action” of increased expression of all genes in the CNV, but from a few critical genes that are deleterious when overexpressed (Bonney et al. 2015).

### CNV strains do not exhibit previously described transcriptional signatures of aneuploidy

Previous studies of a laboratory strain of yeast (W303) identified a transcriptomic signature of aneuploidy independent of which chromosome is duplicated (Torres et al. 2007; Terhorst et al. 2020), that is characteristic of the yeast environmental stress response (ESR) (Gasch et al. 2000) and comprises 868 genes. The expression of genes in the ESR are correlated with growth rate (Brauer et al. 2008) and several studies have shown that strains with higher degrees of aneuploidy (i.e. more additional base pairs) exhibit lower growth rates and stronger ESR (Torres et al. 2007; Terhorst et al. 2020). We compared the ESR gene expression profiles in our CNV strains and a previous study which identified the ESR as a response to aneuploidy (Torres et al. 2007) (**Figure 5A, SFig 12, SFig 13**). Surprisingly, we find a significant negative correlation for all strains except ComTrip, with the strength of the anticorrelation decreasing as growth rate increases.

**Figure 5.**
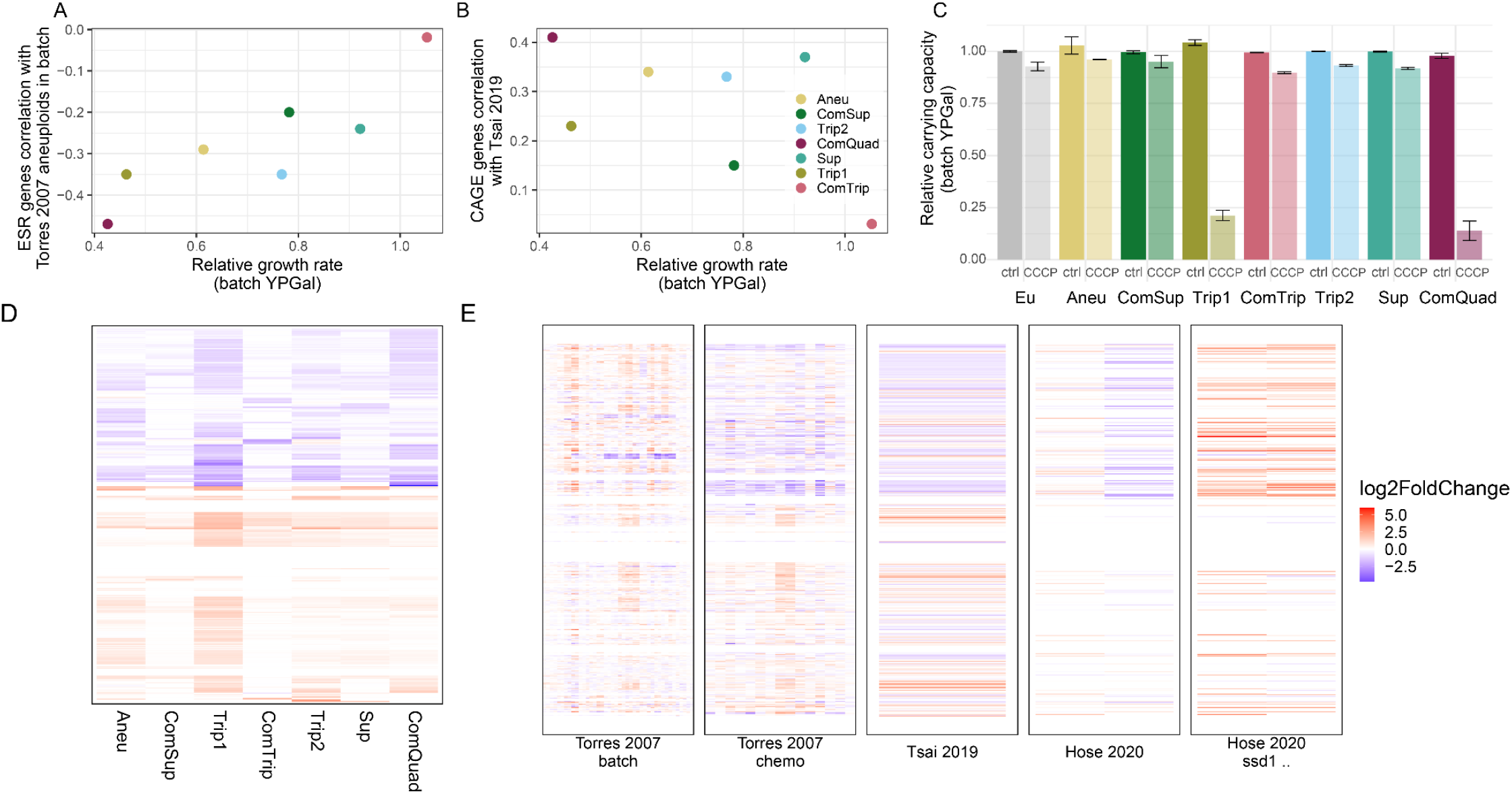
Global gene expression signatures in CNV strains are distinct from aneuploidy effects. We compared the mean growth rate of CNV strains in YPGal to **A)** the Pearson correlation between log2 fold change in mRNA expression in CNV strains vs euploids in this study and the mean log2 fold change in mRNA expression for aneuploids vs euploids grown in batch culture in Torres et al. 2007 for 798 ESR genes for which we have complete data, **B)** the Pearson correlation between log2 fold change in mRNA expression in CNV strains vs euploids in this study and the log2 fold change in mRNA expression in for aneuploids vs euploids in Tsai et al. 2019 for the 215 CAGE genes for which we had complete data. **C)** Average and standard deviation (error bars) carrying capacity (i.e. maximum optical density) relative to the ancestral, euploid strain in YPGal batch culture in either control condition or with 25 μM CCCP. **D)** Log2 mRNA expression for 436 genes (rows) significantly differentially expressed in at least one CNV strain versus the euploid strain. **E)** Data corresponding to genes from **(D)** from Torres et al. 2007 aneuploids in batch, Torres et al. 2007 aneuploids in chemostat, Tsai et al. 2019, Hose et al. 2020 wild aneuploid strains, and Hose et al. wild aneuploid strains with ssd1 deleted. The former four are compared to closely related euploids, the aneuploids with ssd1 deletions are compared to their wild-type aneuploid counterparts.

Recently, a common aneuploidy gene-expression (CAGE) signature across yeast strains aneuploid for many different chromosomes that is similar to the transcriptional response to hypo-osmotic shock was defined in a derivative of the S228C genetic background (Tsai et al. 2019). With the exception of ComTrip, expression of CAGE genes is moderately positively correlated between Tsai et al. (2019) and our CNV strains (**Figure 5B, SFig 14**). Interestingly, expression of CAGE genes is also positively correlated with expression of CAGE genes in growth rate controlled aneuploid strains in Tsai et al. (2019) (S**Fig 15**).

### Genome-wide gene expression effects of CNVs

From our differential analysis, we identified 436 genes, 341 of which are not located on chromosome XI, that had significantly altered expression in one or more CNV strains compared to the euploid strain (log2 fold change > 1.5, Benjamini and Hochberg adjusted p < 0.05, **Figure 5D, STable 23**). Of the significant genes, 73 are ESR genes and 13 are in the CAGE signature. The 436 genes fall into two major clusters; genes that have decreased expression are involved in cellular respiration, nucleoside biosynthetic processes, and small molecule metabolism, and genes that have increased expression are involved in transposition, nucleic acid metabolic processes, and siderophore transport (hypergeometric test p < 0.0001, **STable 24**). Similarly, wild yeast strains that are aneuploidy tolerant exhibit down-regulation of mitochondrial ribosomal proteins and genes involved in respiration, and upregulation of oxidoreductases (Hose et al. 2015). The clusters and enrichment patterns remain even when excluding genes on chromosome XI (**STable 25**). Additionally, we see similar functional enrichment for gene expression differences of individual strains (**STable 18**).

There are nine genes that have significantly different expression than the euploid in all strains and are not on chromosome XI (S**Fig 16**). Three genes have increased expression: two are retrotransposons and the third, *RGI2*, is a protein of unknown function that is involved in energy metabolism under respiratory conditions (Domitrovic et al. 2010). Repressed genes include the paralogs *MRH1* and *YRO2*, both of which localize to the mitochondria (Reinders et al. 2007, 2006); *OPT2*, an oligopeptide transporter (Wiles et al. 2006); *YGP1*, a cell wall-related secretory glycoprotein (Destruelle et al. 1994); and two proteins of unknown function, *PNS1* and *RTC3*.

We compared the genes that are significantly differentially expressed in one or more CNV strains to the data generated from aneuploid strains in (Torres et al. 2007; Tsai et al. 2019; Hose et al. 2020) (**Figure 5E**). As with comparison to the ESR and CAGE signatures, we see that our strains are more similar to the aneuploids grown in chemostats (Torres et al. 2007) and the aneuploids (Tsai et al. 2019) than the aneuploids growing in batch culture and exhibiting the ESR. Hose (Hose et al. 2020) compared gene expression in aneuploid wild yeast strains that are tolerant of CNV to their euploid counterparts and to aneuploid wild yeast with *SSD1* deleted. The gene expression profiles of the aneuploid wild yeast more closely resemble our strains than the aneuploid *SSD1* mutants, which are more similar to the aneuploids grown in batch culture in (Torres et al. 2007) (**Figure 5E**).

### Low fitness is associated with mitochondrial dysfunction

Several lines of evidence suggested that the slower relative growth rates of strains Aneu, Trip1, and ComQuad (**Figure 3D**) may be the product of mitochondrial dysfunction. Not only did these strains have heightened tolerance for *BMH1* insertions (**Figure 3A**), which exhibits positive genetic interactions in most CNV strains (**Figure 3E**), BMH1 is also a negative regulator of retrograde signaling (da Cunha et al. 2015). These strains also exhibited shared genetic interactions and mRNA expression signatures linked to mitochondrial function and translation activity (da Cunha et al. 2015). One possible explanation of these signatures is that the mitochondria are dysfunctional and therefore retrograde signaling is constitutively activated, allowing a relaxation of insertions in *BMH1*. However, *CIT2* and *DLD3*, which are robustly upregulated in the canonical retrograde response (da Cunha et al. 2015), do not have significantly different expression from the euploid in any CNV strain.

Yeast simultaneously ferment and respire galactose (Fendt and Sauer 2010). Therefore, to test whether CNV lineages have impaired mitochondrial function, we tested growth in the presence of carbonyl-cyanide 3-chlorophenylhydrazone (CCCP), a mitochondrial uncoupling agent. We found that treating with CCCP nearly abolished growth in two (Trip1, and ComQuad) of the three strains that showed strongest signals differential mitochondrial function, whereas the reduction in growth in other strains was similar to that of the euploid (**Figure 5C**). Looking at expression patterns unique to these strains we found a variety of gene sets enriched in mitochondria associated pathways and functions (**SFig 17**) These results are consistent with impaired mitochondrial function in these strains, as suggested by the previously described *BMH1* insertion tolerance and the reduced growth on glycerol phenotype.

## Discussion

In this study, we sought to understand the effect of diverse CNVs on genetic interactions and gene expression. Though investigations of evolutionary trajectories and combinations of mutations that arise in evolution experiments have suggested that epistasis between CNVs and other mutations is an important contributor to evolutionary dynamics (Pavani et al. 2021; Lauer et al. 2018), few studies have systematically investigated genetic interactions with CNVs (Dodgson et al. 2016). We find that gene amplification results in relaxed selection against mutation in essential genes. Additionally, we find both CNV specific genetic interactions and interactions that are shared by several strains. To gain further insight into the effects of CNVs on the cells, we performed gene expression analysis using RNAseq. Whereas gene amplification results in increased mRNA expression compared to the euploid, we do not find evidence for dosage compensation as gene expression increases are explained by increased template number. Consistent with a recent study in the same genetic background as our strains (S288C), we do not find activation of the ESR in various aneuploids (Larrimore et al. 2020), nor did we observe the CAGE response (Tsai et al. 2019). Instead, CNVs exhibit a unique gene expression signature comprising increased expression of genes involved in transposition, nucleic acid metabolic processes, and siderophore transport, and decreased expression of those involved in cellular respiration, nucleoside biosynthetic processes, and small molecule metabolism, though the extent to which the expression differed from the euploid varied between CNV strains.

We have demonstrated that transposon mutagenesis is a powerful tool to investigate genetic interactions genome-wide in strains with large and complex mutations. Unlike synthetic genetic array (SGA) analysis, which is commonly used to investigate genetic interactions, transposon mutagenesis does not require mating the query strain to the deletion collection. Transposon mutagenesis therefore avoids a some of the issues that are encountered using SGA: inaccuracies in the deletion collection (Giaever and Nislow 2014; Ben-Shitrit et al. 2012), secondary mutations including aneuploidy (Hughes et al. 2000), and the gene expression of non-target genes is sometimes impacted by the deletion of neighboring genes (Ben-Shitrit et al. 2012; Baryshnikova and Andrews 2012) which can result in false positive or false negative genetic interactions. Furthermore, the requirement to mate the query strain to the deletion collection means that genetic interactions identified are in a diploid and potentially hybrid (if the query strain and deletion collection are different) background, which further complicates the interpretation of results (Dodgson et al. 2016). Despite overcoming some of these challenges, transposon mutagenesis also has some shortcomings. Transposon insertion efficiency can differ between genetic backgrounds (Caudal et al.) and ability to detect interactions is dependent on the number of insertions identified. Our experimental design induces transposition in galactose for four days, which may be enough time for additional mutations to also accumulate, reducing the number of conditions in which genetic interactions can be examined and power of the experiment to detect loci which tolerate mutation.

A question that naturally arises from our study is why do different CNV structures result in heterogeneous fitness effects, genetic interactions, and transcriptional responses? One reason may be the particular composition of the CNV. Both the copy number and the particular genes amplified likely play a role in the fitness effect. For example, the aneuploid was much less fit in YPGal than several of the other CNV strains. Previous work has shown that the amplification of the left arm of chromosome XI has negative fitness effects in other conditions (Sunshine et al. 2015) - the aneuploid is the only strain with the left arm amplified in our experiment, and that could be part of the basis of the fitness consequences. Additionally, CNV strains used here were each isolated from an evolution experiment, and other mutations are present in each strain. Given the numerous genes involved in CNVs and the deeply interconnected yeast gene regulatory network, it is reasonable to assume these mutations act in concert with CNVs For example, ComTrip, which has fitness, genetic interactions, and transcriptome similar to the ancestor in YPGal, has a mutation causing a premature stop in *SSK2* (**STable 1**). *SSK2* is a MAP kinase kinase kinase of *HOG1* signaling pathway that controls osmoregulation, which may attenuate stress from CNV if it is, as Tsai et al. (2019) found in aneuploids, similar to hypo-osmotic stress. Further studies in CNVs of various structures encompassing different regions and in isogenic backgrounds may help to disentangle these factors.

Using long-read sequencing, in addition to short-read (Illumina) whole genome sequencing, we were able to not only reconstruct the CNV topologies but also accurately measure copy number change of rDNA locus in the adapted CNV strains (**STable 3**). While rDNA locus copy number change has previously been implicated in response to stress (Salim et al. 2017; Matos-Perdomo and Machín 2019) it is rarely included in the analysis of aneuploidy and CNVs because of the difficulty in accurately measuring the copy number. Interestingly, we find both amplifications and reductions of the rDNA locus in the CNV containing strains, suggesting that the fitness effect of rDNA copy-number may be dependent on the genetic background.

A large-scale analysis of aneuploidy across over 1,000 *S. cerevisiae* isolates showed that genetic background alone (rather than ecology) could predict aneuploidy prevalence (Scopel et al. 2021), and several studies have shown that tolerance to aneuploidy varies across genetic backgrounds (Hose et al. 2015; Gasch et al. 2016; Hose et al. 2020; Larrimore et al. 2020). This understanding leads us to important questions: is there a common response to aneuploidy and more generally the presence of CNVs in those genetic backgrounds that do not well tolerate them? Can we predict which genetic backgrounds will be able to tolerate CNVs or will be sensitive to them? Hose et al. found that aneuploidy sensitivity in the laboratory strain W303 resulted from synergistic defects in mitochondrial function and proteostasis. Interestingly, our results also point to mitochondrial function and translation as important and the least fit strains in our study are particularly sensitive to mitochondrial stress. The laboratory strain S288C, which was used in this study, has a hypomorphic allele of *HAP1 (Gaisne et al. 1999*), which is a heme-responsive transcriptional activator of genes involved in respiration (Zhang and Hach 1999) (notably, genes with increased expression in CNV strains include siderophores, which chelate iron). Across many genetically distinct strains of yeast, genes involved in aerobic respiration and the electron transport chain vary more than any other category during growth in glucose-limited chemostats (Skelly et al. 2013), and genes involved in mitochondrial function have continuous variation in fitness effects across different isolates (Caudal et al. 2022). It would be instructive to study the relationship between mitochondrial function and CNV tolerance.

## Methods

### Yeast Strains

The euploid ancestral *GAP1* CNV reporter and the evolved *GAP1* CNV strains were previously described and characterized in Lauer et al. 2018, and are haploid derivatives of the reference strain S288C (and more specifically, FY4/5) with a constitutively expressed *mCitrine* gene and KanMX G418-resistance cassette inserted 1,118 base pairs upstream of *GAP1*. This construct is referred to as the *GAP1* CNV reporter. The CNV strains are clonal isolates that evolved for 150 or 250 generations in glutamine limited chemostats (Lauer et al. 2018).

Each strain was transformed with pSG36_HygMX using the EZ-Yeast™ Transformation Kit (MP Biomedicals, cat #2100200). Transformants were recovered on YPG agar + 200 μg/mL Hygromycin B. A single colony was picked from the plate of transformants to perform each transposon mutagenesis experiment. Separate transformation and colony selection was performed for each replicate of the euploid.

To generate *BMH1* mutants, we performed high-efficiency yeast transformation into frozen competent yeast cells for each strain (Gietz and Schiestl 2007) with an *mCherry* gene under control of the constitutively expressed *ACT1* promoter (*ACT1pr::mCherry::ADH1term*) and marked by the HphMX Hygromycin B-resistance cassette (*TEFpr::HygR::TEFterm*). The plasmid DGP363, containing this construct, was used as template for PCR using primers containing the same *BMH1-specific* targeting homology, and transformation resulted in a complete deletion of the *BMH1* open reading frame. Transformants were recovered on YPD agar + 400 μg/mL G418 + 200 μg/mL Hygromycin B, and *BMH1* deletion positive transformants were confirmed using *BMH1* specific primers and a HygR primer. We verified that *mCitrine* copy number remained unchanged and *mCherry* fluorescence using a Cytek Aurora flow cytometer.

### Growth analysis

For each experiment, we inoculated three colonies per strain into 3-5 mL YPGal, and grew them overnight at 30°C. In triplicate per original colony, we back diluted 5 μL of culture into 195 μL fresh YPGal or YPGal with 25 μM carbonyl-cyanide 3-chlorophenylhydrazone in a Costar Round Bottom 96 well plate (Ref 3788). We treated the lid with 0.05% Triton X-100 in 20% ethanol to prevent condensation (Brewster 2003). We collected OD600 data over approximately 48 hours using a Tecan Spark with the following parameters: Temperature control: On; Target temperature: 30 [°C]; Kinetic Loop; Kinetic cycles: 530; Interval time: Not defined; Mode: Shaking; Shaking (Double Orbital) Duration: 240 [s]; Shaking (Double Orbital) Position: Current; Shaking (Double Orbital) Amplitude: 2 [mm]; Shaking (Double Orbital) Frequency: 150 [rpm]; Mode: Absorbance; Measurement wavelength: 600 [nm]; Number of flashes: 10; Settle time: 50 [ms]; Mode: Fluorescence Top Reading; Excitation: Monochromator; Excitation wavelength: 497 [nm]; ExcitationBandwidth: 30 [nm]; Gain: Calculated From: B5 (50%); Mirror: AUTOMATIC; Number of flashes: 30; Integration Time: 40 [μs]; Lag time: 0 [μs]; Settle time: 0 [μs]; Z-Position mode: From well B5.

Using growthcurver (Sprouffske and Wagner 2016), we fit the OD600 data to a logistic equation, using the value of the parameter *r* as the intrinsic growth rate of the population and the parameter *k* as the carrying capacity. We checked for and discarded outliers by examining OD curves and histograms of sigma (goodness of fit). We normalized each growth rate and carrying capacity to that of the ancestral euploid *GAP1* CNV reporter grown in the same plate.

### Transposon mutagenesis

A single transformant for each strain was used to inoculate a 30 mL YPD + 200 μg/mL Hygromycin B, and incubated approximately 24 hours at 30°C with agitation, until OD5. To induce transposition, the culture was then diluted to OD0.05 in YPGalactose + 200 μg/mL Hygromycin B to a final volume of 50 mL, and incubated 24 hours at 30°C with agitation. The culture was diluted to 0.05 in 50 mL YPGalactose + 200 μg/mL Hygromycin B and incubate 24 hours three more times, for a total of four serial transfers in YPGalactose + 200 μg/mL Hygromycin B. The culture was pelleted by centrifugation for five minutes at 4000 rpm, the supernatant removed, then resuspended to OD0.5 in 50 mL YPD and incubated 24 hours at 30°C with agitation, then diluted again to OD0.5 in 50 mL YPD and incubated 24 hours at 30°C with agitation, to release selection to maintain pSG36_HygMX. The cultures were then diluted to OD0.5 in 100 mL YPD + 200 μg/mL Hygromycin B and incubated 24 hours at 30°C with agitation to select for cells with the transposon in the genome. The final culture was pelleted by centrifugation for five minutes at 4000 rpm, the supernatant removed, resuspended with 1 mL sterile water, split into four 250 μL aliquots, and pelleted for two minutes at 8000 rpm. The supernatant was removed and cell pellets were frozen at −20°C for storage until DNA extraction was performed.

### Insertion site sequencing

DNA was extracted from cell pellets using the MasterPure™ Yeast DNA Purification Kit (Lucigen, cat #MPY80200), with an additional initial incubation with zymolyase at 37°C to enhance cell lysis, and using a Glycogen/Sodium Acetate/Ethanol DNA precipitation(Green and Sambrook 2016). For each sample, 2 μg of DNA was digested with 50 units of DpnII and 5 μL NEBuffer™ DpnII (NEB, cat #R0543L), in a total volume of 50 μL; and 2 μg of DNA was digested with 50 units of NlaIII and 5 μL CutSmart^®^ Buffer (NEB, cat #R0125L), in a total volume of 50 μL, for 16 hours at 37°C. The reactions were heat inactivated, then circularized by ligation in the same tube with 25 Weiss units T4 Ligase and 40 μL T4 ligase buffer (Thermo Scientific cat #EL0011) for 6 hr at 22°C, in a volume of 400 μL. Circularized DNA was precipitated using a Glycogen/Sodium Acetate/Ethanol DNA precipitation (Green and Sambrook 2016). Inverse PCRs for each sample and digestion were performed with primers Hermes_F and Hermes_R with 0.5 μL of each circularized DNA sample per reaction. PCR was performed with DreamTaq (ThermoFisher cat #EP0701), with the following program: 2 min at 95°C followed by 32 cycles of 30 s at 95°C, 30 s at 57.6°C, 3 min at 72°C, and a final extension step of 10 min at 72°C. The PCRs products were confirmed on 2% agarose gels, and the concentration was quantified using Qubit™ dsDNA BR Assay Kit.

Library preparation and sequencing were performed using two different library preparation methods and sequencing set ups as follows

#### BGI

For each sample (1728, 1736, and 1740) and digestion, 35 PCR reactions using primers (**STable X10**) each with 0.5 μL of each circularized DNA were performed as described above and the PCR products were pooled and cleaned using a Glycogen/Sodium Acetate/Ethanol DNA precipitation (Green and Sambrook 2016). For each sample, at least 6 μg at minimum 30 ng/μl was then sent to the BGI (Beijing Genomics Institute) for library preparation and sequenced using a paired-end (2 x 100) protocol on a Illumina Hi-Seq 4000 or DNBseq platform.

#### NYC

For each sample (all) and digestion 4 PCR reactions were performed as described above and the PCR products were pooled by sample and cleaned using a Glycogen/Sodium Acetate/Ethanol DNA precipitation (Green and Sambrook 2016). Five ng of each PCR product pool was used as input into a modified Nextera XT library preparation. To increase library complexity, for each sample, two tagmentation reactions were performed. PCR to enrich for fragments with hermes sequence and add an i5 adaptor were performed on the tagmented DNA using NPM Buffer, primers Nextera_hermes_enrichment and Nextera_i7_enrichment, and the following program: 3 min at 72°C, then 30 s at 95°C, followed by 9 cycles of 10 s at 95°C, 30 s at 55°C, 30 s at 72°C, and a final extension step of 5 min at 72°C. The reactions were pooled by sample, cleaned using AmPure XP beads, and resuspended in 20 μL of molecular grade water, which was used as input for an indexing and library amplification PCR. Each sample was indexed with an i7 index from the Nextera XT kit, and amplification of the i5 end was performed with primer i5_amp (which contains no i5 index), using the 2X KAPA PCR master mix (Roche cat. #KK2611), and the same program described for the PCR after tagmentation.PCR cleanup and size selection was performed with AmPure XP beads. The fragment size of each library was measured with an Agilent TapeStation 2200 and qPCR was performed to determine the library concentration. The libraries were pooled at equimolar concentrations, and sequenced using a single-end (1 x 150) protocol on an Illumina NextSeq 500. Libraries were prepared once, but sequenced in two consecutive sequencing runs for increased coverage.

### Transposon insertion sequencing site identification and annotation

Using cutadapt v1.16 (Martin 2011) with the expected hermes TIR sequence on the 5’ end were identified, and the TIR was trimmed. If the TIR was followed by plasmid sequence, these reads were discarded. For reads sequenced at BGI (paired end sequencing), the read with the TIR sequence was identified and its mate was discarded. For reads sequenced at NYC (Nextera based prep, single end sequencing), Nextera transposase sequences were identified and removed. Reads with a length less than 20 bases after all cleaning steps were discarded, and the remaining reads were checked for quality using fastqc v0.11.8 ([CSL STYLE ERROR: reference with no printed form.]). Reads were aligned to the modified reference genome using bwa mem v.0.7.15 (Li and Durbin 2010) and BAMs were generated with samtools v1.9 (Li et al. 2009). Samples prepared and sequenced by more than one method had high Pearson correlations (0.85-0.94) in the number of unique insertions identified per gene (**STable X3-pre12**), and therefore were combined into a single BAM file before performing downstream analysis. For the majority of the analyses, BAMs were combined by sample, for ease of processing and to prevent redundant insertion site identification. BAMs were parsed with a custom python script which identifies the first base of the read as the position of the insertion. The script output all unique insertion positions and the number of reads per insertion position. Positions were annotated using bedtools v2.26.0 (Quinlan and Hall 2010) and a custom GFF containing amended annotations for the custom genome (Supplemental File X1 - GAP1.gff). All analyses use unique insertion positions, and do not take into account the number of reads per unique insertion position. Uniquely identified insertion sites are supported by an average of 18.6 sequencing reads. The libraries have between 85,327 and 329,624 unique insertion sites identified, with an average of 176,664 insertion sites, corresponding to approximately one insertion per 69 bases in the yeast genome (NCBI R64 assembly; **STable 2**). We normalize for differences in sequencing depth by calculating insertions per million: number of unique insertion sites per feature/(total unique insertion sites/1,000,000) (Levitan et al. 2020). We do not normalize for gene length, as we are comparing genes between strains, not within strains. All code used for analysis can be found on GitHub https://github.com/graceave/hermes_analysis.

### Genetic interaction analysis

To quantitatively investigate genetic interactions using the transposon sequencing data, we performed differential analysis using DESeq2 version 1.30.1 (Love et al. 2014), using the number of insertions per gene and comparing each CNV strain to the two euploid replicates. We used clusterProfiler version 3.18.1 (Yu et al. 2012) to perform fast gene set enrichment analysis (Korotkevich et al.) using the ranked log2 fold change in insertions generated by DESeq2 and GO terms were summarized by semantic similarity (Yu et al. 2010) then by hand for clarity (**STable 4**).

To calculate genetic interactions based on growth rates, we first calculate the relative fitness of each single mutant by:

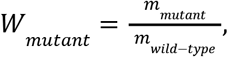

where *m* is the intrinsic growth rate of the strain (parameter *r* from logistic equation used to fit growth curves). We then calculated the expected fitness of the double mutant using either the additive model:

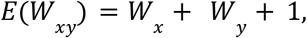

Or the multiplicative model:

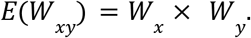

We then calculate the genetic interaction:

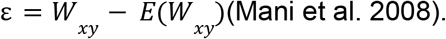

### RNA sequencing

For RNA sequencing, we grew overnight cultures from three replicate colonies per strain in 5 mL YPGal, then 2 mL (euploid, ComTrip) or 5 mL (other strains) of overnight culture was pelleted and subsequently resuspended in 5 mL fresh YPGal. The cultures were allowed to grow for three hours in fresh YPGal before harvesting cells by vacuum filtration and fixing immediately in liquid nitrogen, so that all cultures were harvested while cells were proliferating. RNA was extracted and purified using a hot acid phenol/chloroform and Phase Lock Gels as described in (Neymotin et al. 2014). Samples were enriched for polyadenylated RNA using the Lexogen Poly(A) RNA Selection Kit V1.5 (cat. # 157.96) and stranded RNAseq libraries were prepared using the Lexogen CORALL Total RNA-Seq Library Prep Kit (cat. # 095.96) according to the manufacturer’s protocol. The libraries were pooled at equimolar concentrations, and sequenced using a paired-end (2 x 150) protocol on an Illumina NextSeq 500. The resulting FASTQs were trimmed, aligned, and UMI deduplicated, and coverage per feature was calculated using an in-house pipeline which can be found at https://greshamlab.bio.nyu.edu/wp-content/uploads/2021/11/Windchime_pipeline.nb_.html. Coverage per feature correlation between replicates was high, with the exception of one replicate of ComQuad, which was excluded from further analysis (**STable X3-pre13**). Trip1 and ComTrip also only had two replicates, as library preparation failed for one replicate in each.

### Gene copy number determination and transcript abundance copy number correction

The determination of copy number for each gene in each strain (**STable 4)** was performed using the reconstructed CNV topologies using hybrid long-read and short read sequencing (Spealman et al. 2022).

These copy numbers were then used to make an expected mRNA abundance estimate, or copy number corrected estimate. In order to evaluate dosage compensation of CNVs we sought to have an accurate null model. This expected expression model assumes no dosage compensation, and as such, the expected expression of a CNV associated gene would be equal to the euploid expression multiplied by however many copies of the gene are present in any given strain (**STable 20)**. The difference between the observed and expected expression can then be evaluated using DESeq2 (**STable 21)**, as described above. In the event of CNV dosage compensation one would expect the observed value to be significantly less than the expected value.

## Data Access

All raw and processed sequencing data generated in this study have been submitted to NCBI Sequence Read Archive (SRA; https://www.ncbi.nlm.nih.gov/sra/) under accession number: PRJNA910831.

All code used in computational analysis is publicly available on github: CNV_essentiality

## Supplement

**Supplemental_Fig_S1.**
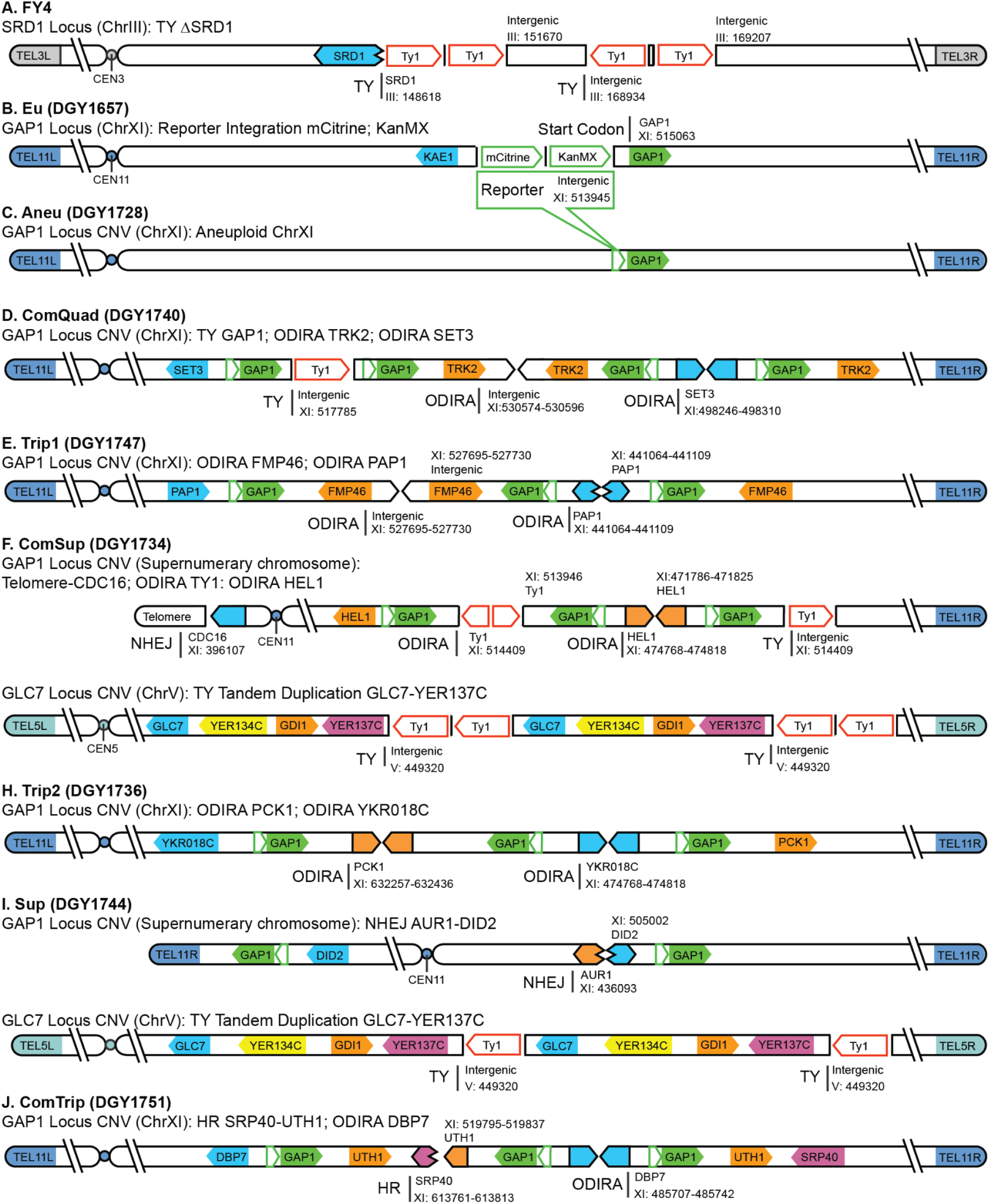
Changes compared to reference S288C genome. Diagrams show the transposon interruption of SRD1 in FY4 (**A**) and the integration of the reporter in the ancestral euploid *GAP1* CNV reporter strain (**B**). Topology diagrams for evolved strains indicating CNV breakpoints, orientations, and the occurrence of transposon events (**C-I**) and most likely mechanism of action TY: transposon-yeast event; ODIRA: origin dependent inverse triplication; NHEJ: non-homologous end joining; HR: homologous recombination

**Supplemental_Fig_S2.**
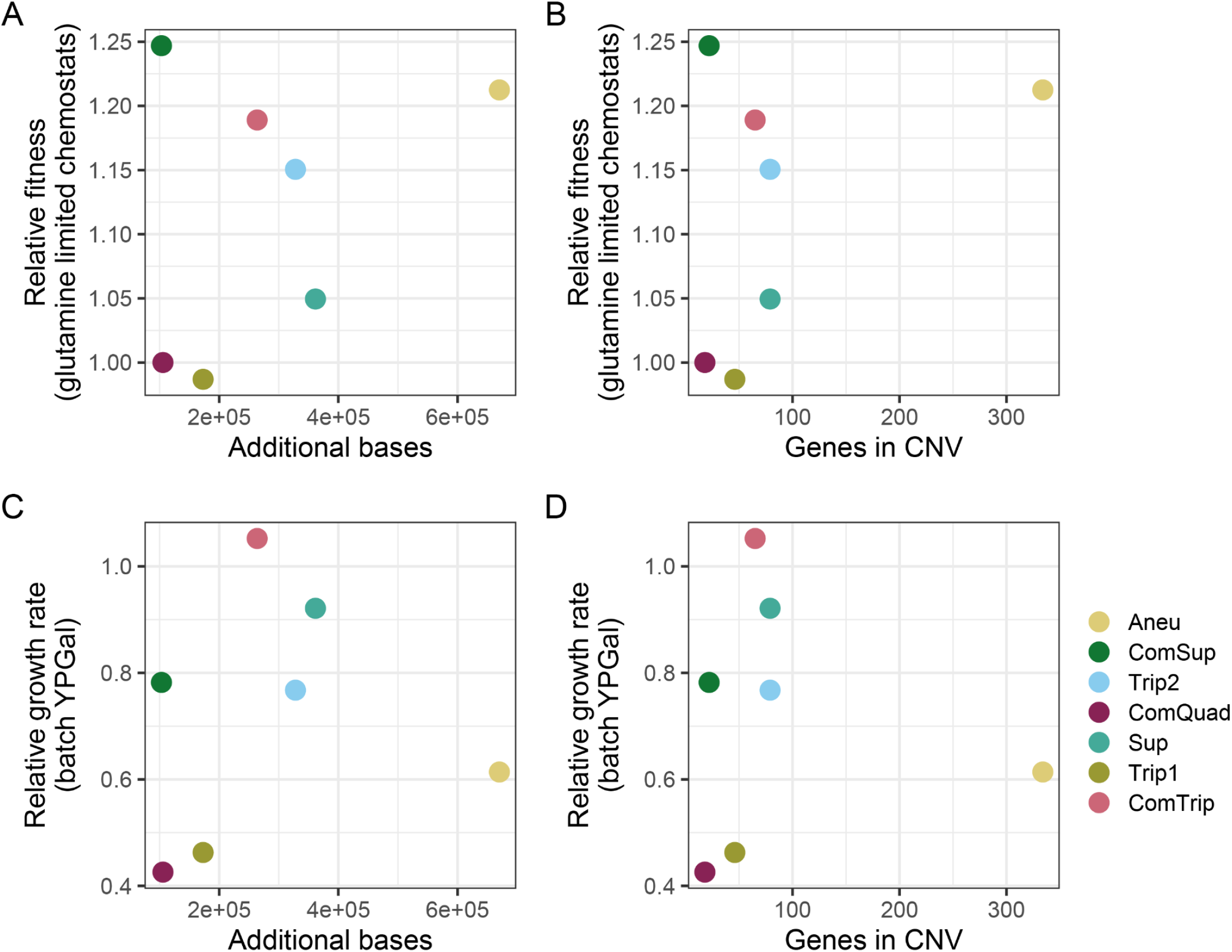
There is no relationship between CNV size and relative fitness. **A-B**) The fitness of evolved strains containing *GAP1* CNVs was determined by pairwise competition experiments with a nonfluorescent, unevolved reference strain in glutamine-limited chemostats. **C-D**) Average growth rate of *GAP1* CNVs relative to the ancestral, euploid strain in YPGal batch culture.

**Supplemental_Fig_S3.**
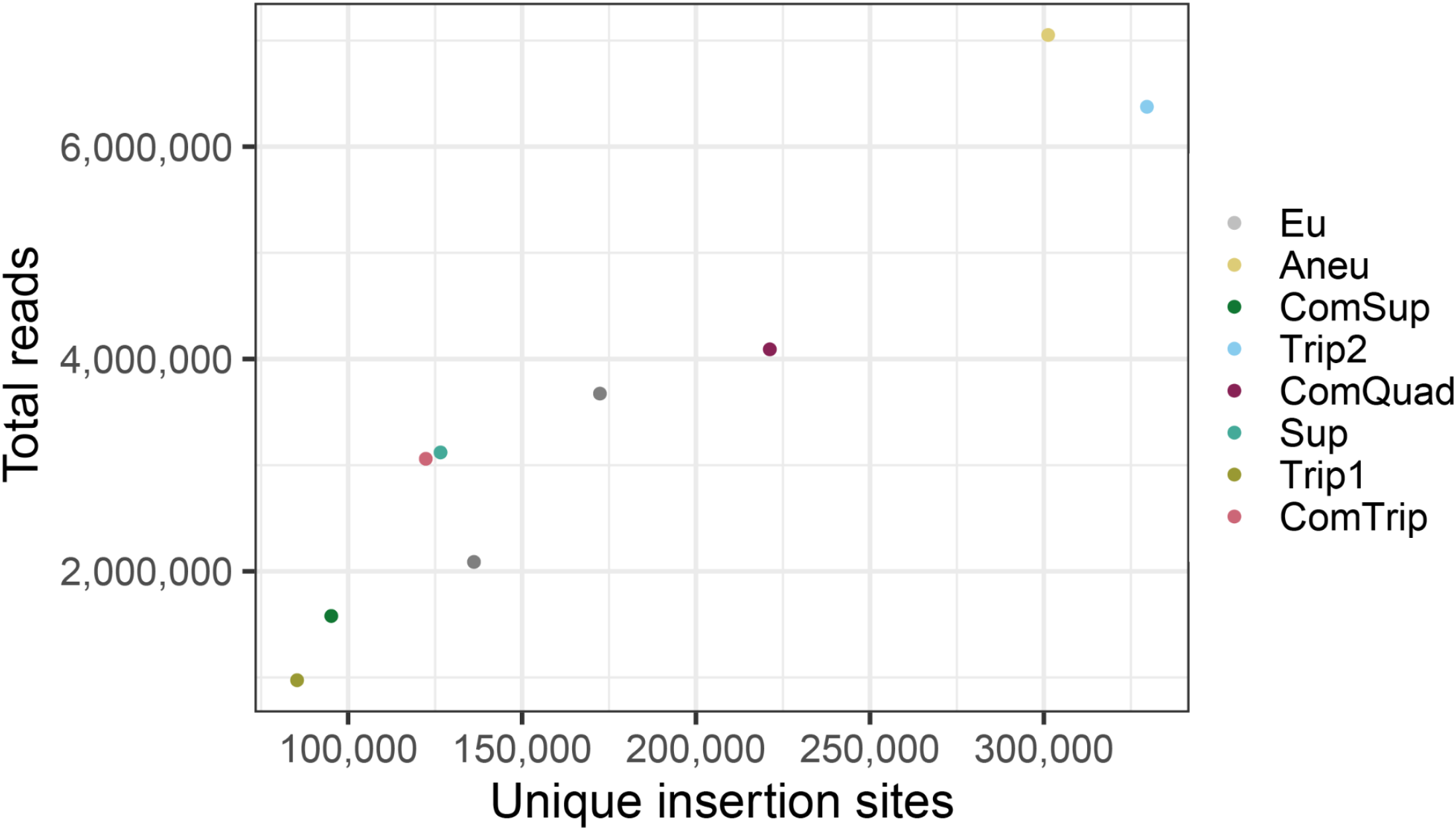
The number of unique insertion sites scales with the number of reads sequenced. The total number of unique insertion sites identified per library increases with the total number of reads sequenced (using all methods and sequencing runs).

**Supplemental_Fig_S4.**
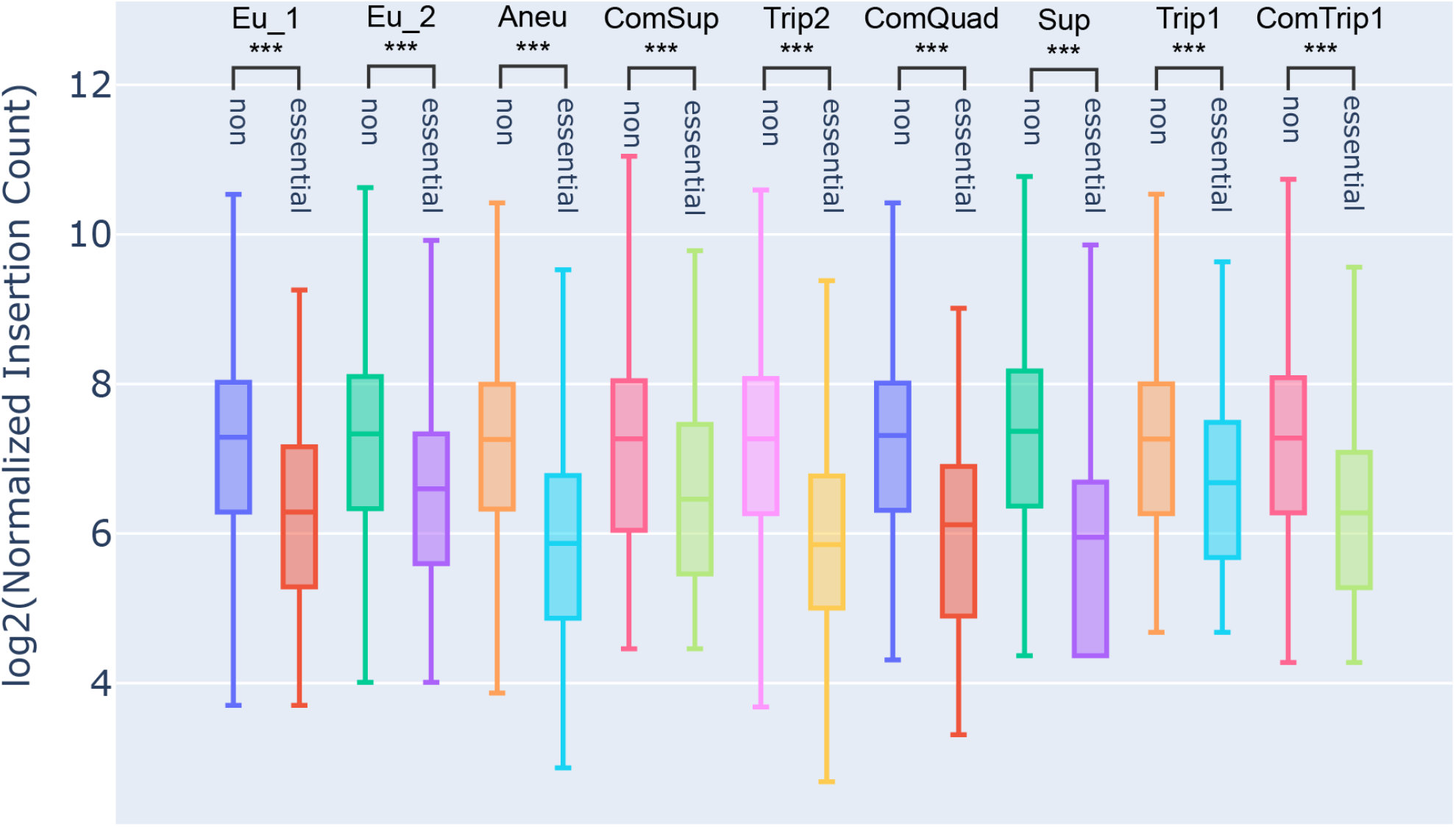
The number of unique insertion sites in the non-essential and essential genes of each strain. We find significantly fewer unique insertions in the essential genes relative to the non-essential genes in each strain (Mann-Whitney U, p-value <= 0.0001).

**Supplemental_Fig_S5.**
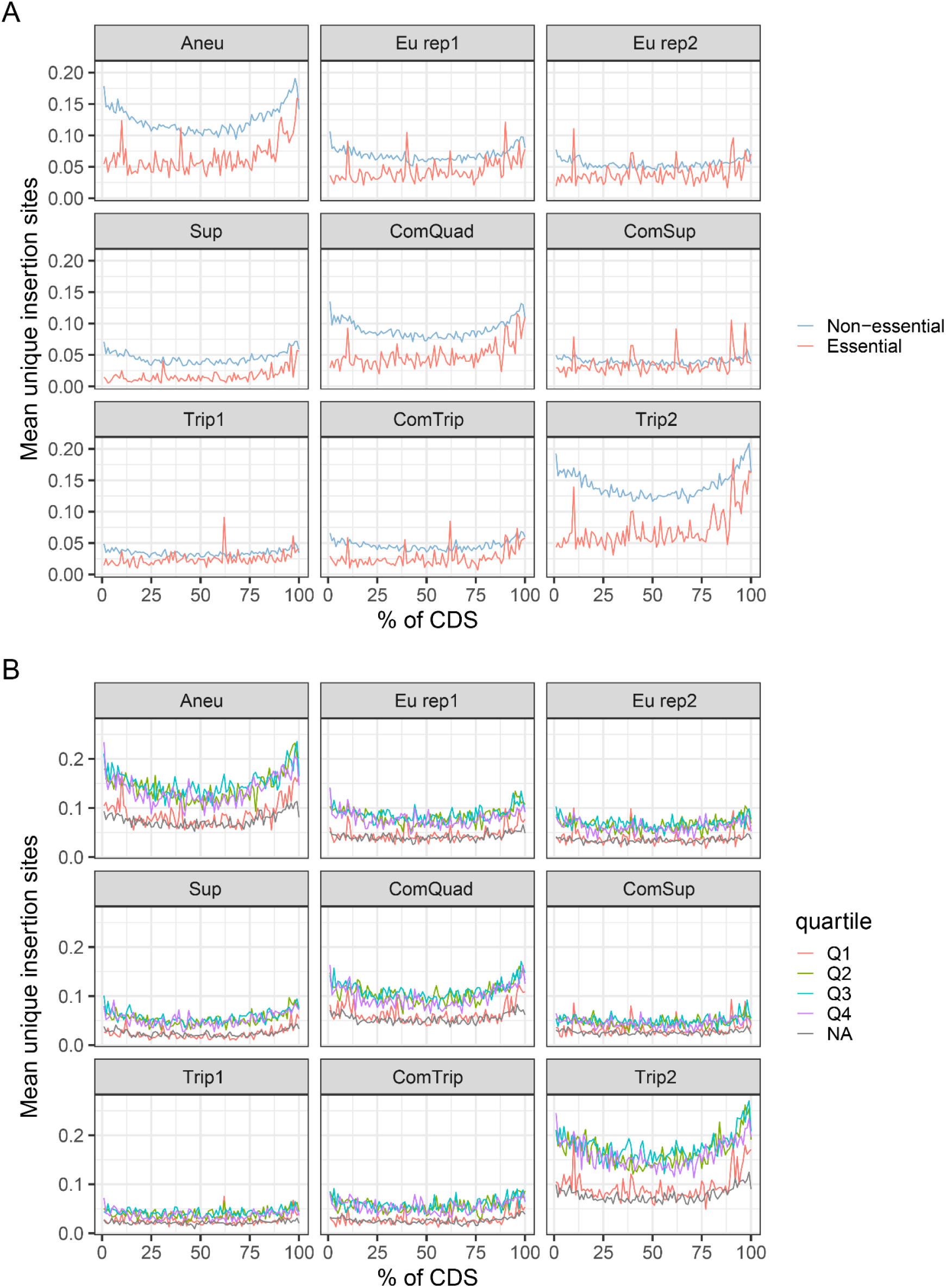
There are fewer insertions in essential genes and genes whose deletion results in low fitness in YPGal. **A)** We grouped genes into those that had been previously annotated as essential or non-essential by deletion and measurement of growth on rich media (yeast peptone dextrose) (Winzeler et al. 1999). **B)** We grouped genes into four quartiles based on relative fitness measurements on rich media with 2% galactose from 3704 viable deletion mutant strains and 782 temperature-sensitive (TS) alleles (Costanzo et al. 2021). The first quartile (Q1, red) contains genes whose deletion causes the greatest measurable fitness defects, with relative fitness between 0.053 and 0.896. There was no relative fitness obtained for 21 genes (presumably there was no growth), these are marked NA (gray).

**Supplemental_Fig_S6.**
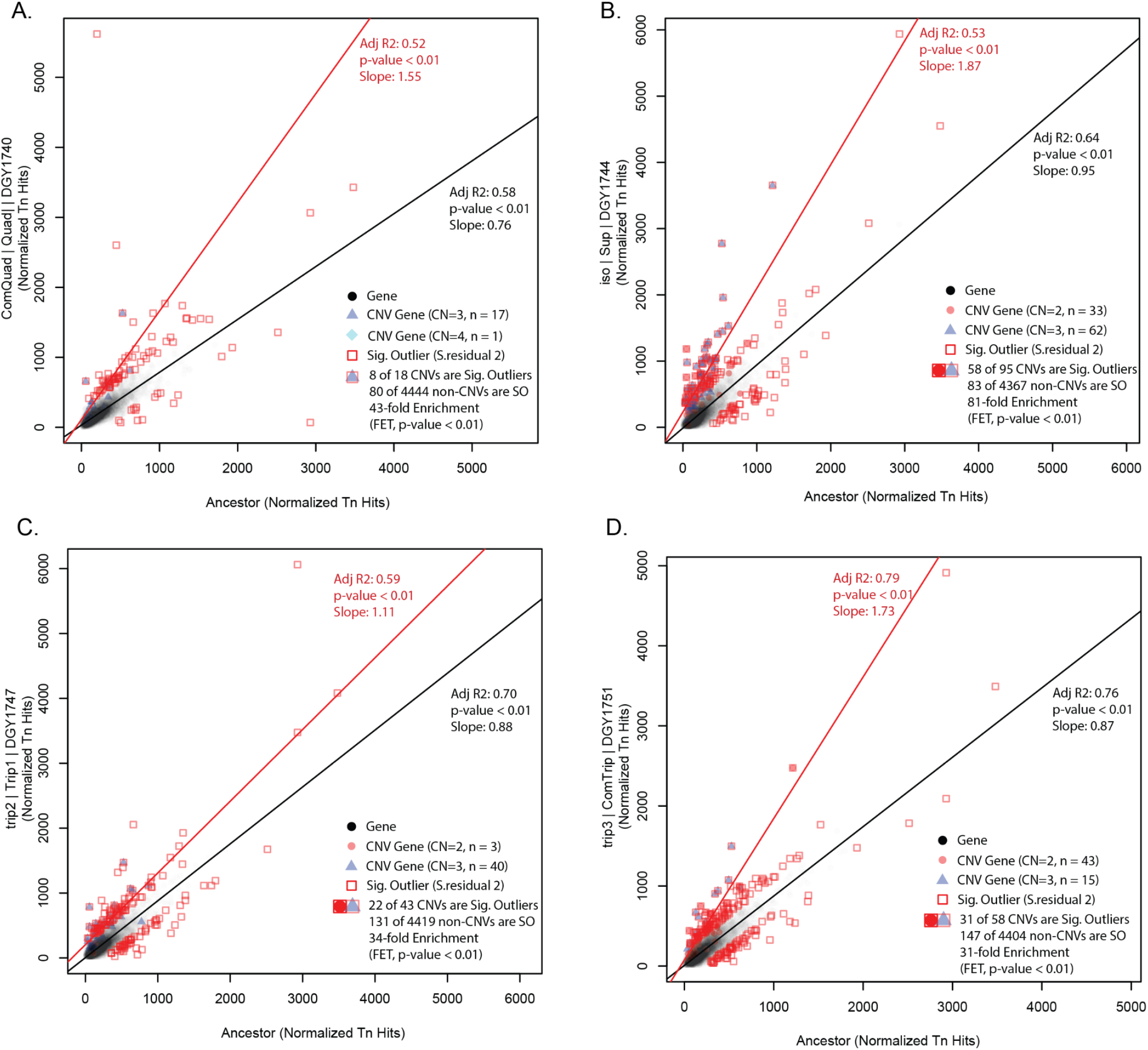
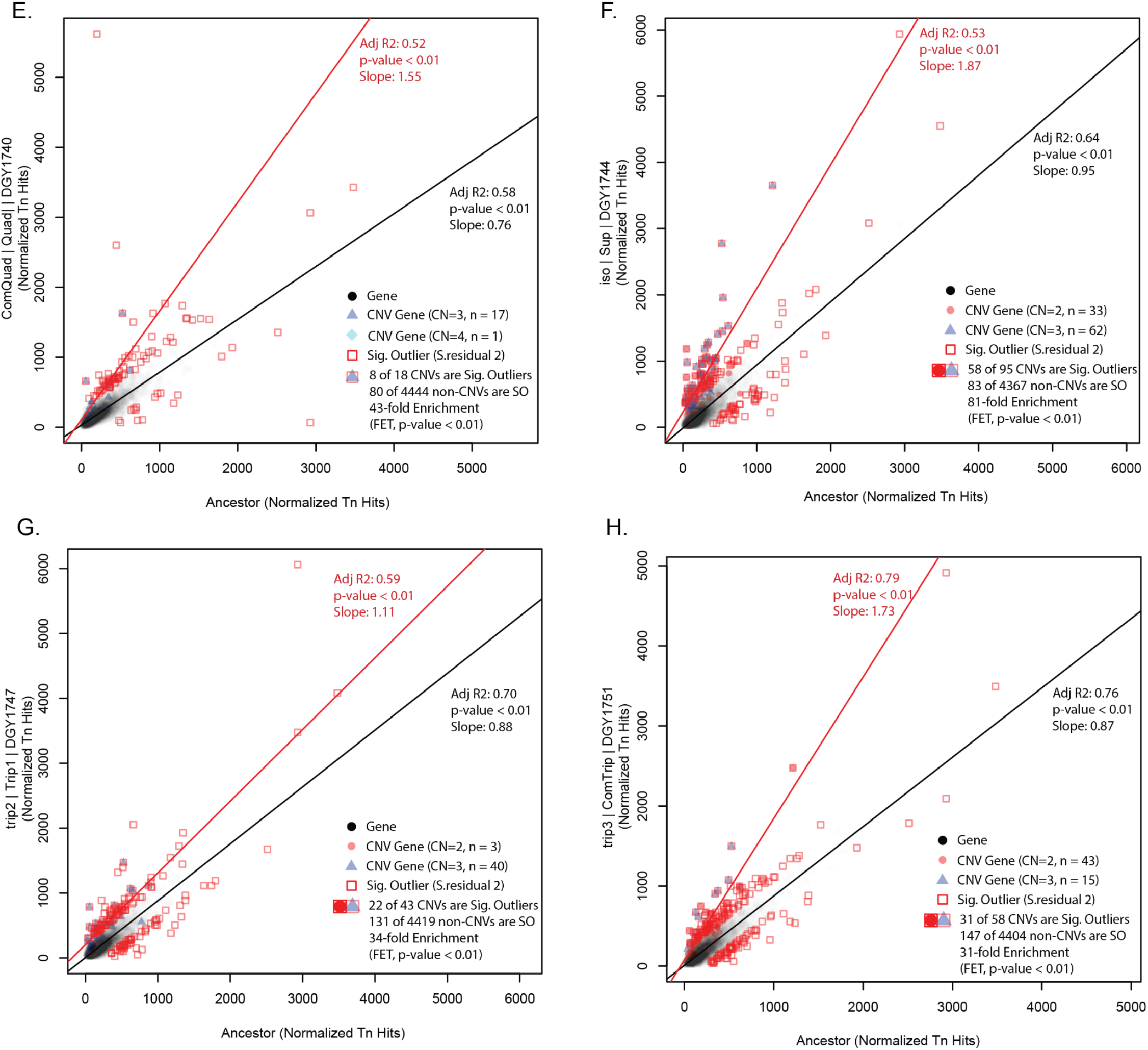
R-squared linear regression of Tn insertions between CNV strains and Euploid ancestor. A R-squared linear regression was performed for all CNV strains relative to the Euploid ancestor. Linear regressions were made for all genes (black circles, black line) or for CNV associated genes (red circles copy number CN= 2, blue triangles CN=3, green diamonds CN = 4, red line). Significant outliers are genes with standardized residuals greater than 2 (red squares). The enrichment of significant outliers in CNV associated genes relative to their occurrence in non-CNV associated genes was calculated using Fisher exact test. In each strain the CNV associated genes were significantly enriched in outliers (FET, p-value < 0.01).

**Supplemental_Fig_S7.**
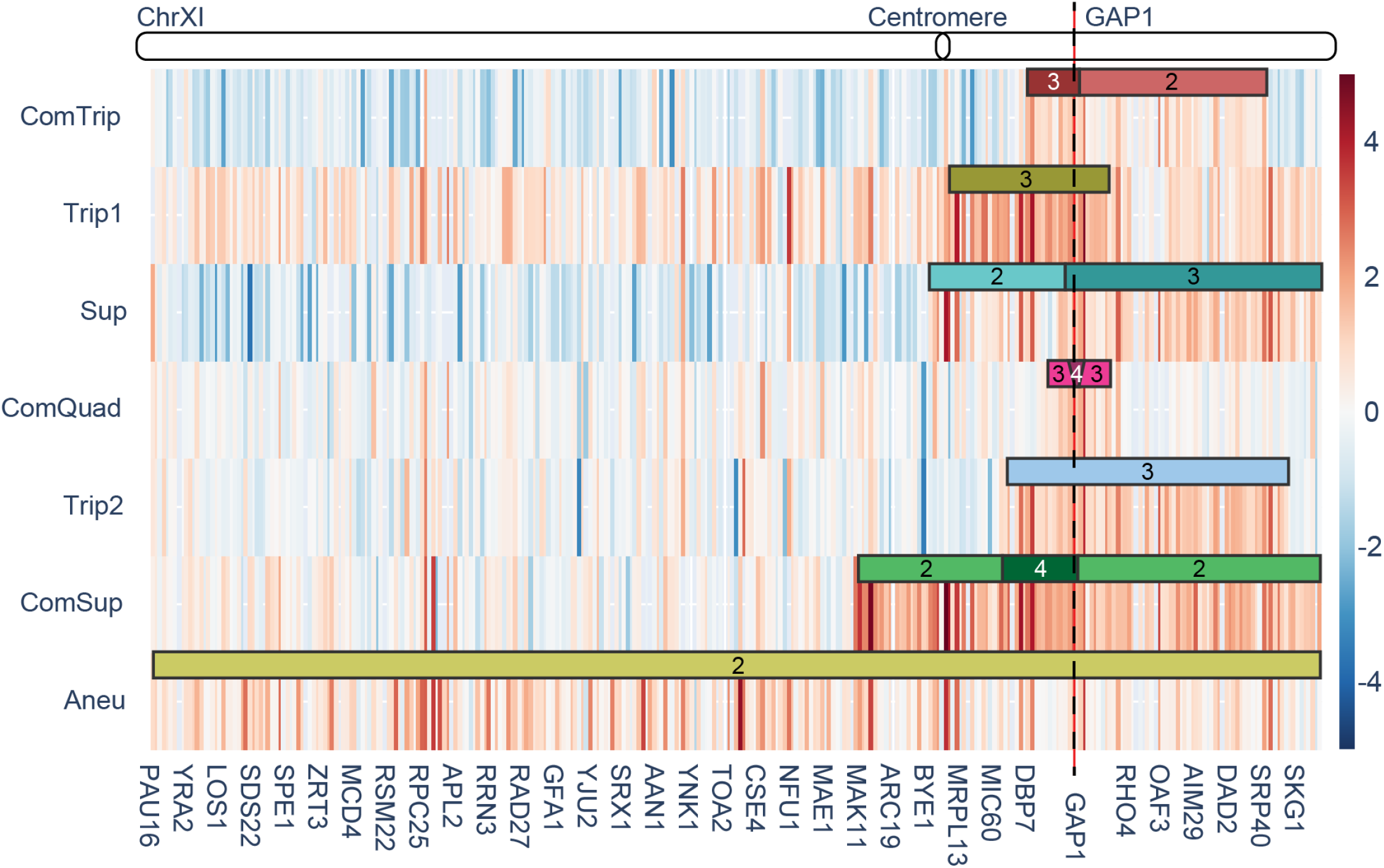
Heatmap of unique transposon insertions counts and Chromosome XI map with CNV regions.

**Supplemental_Fig_S8.**
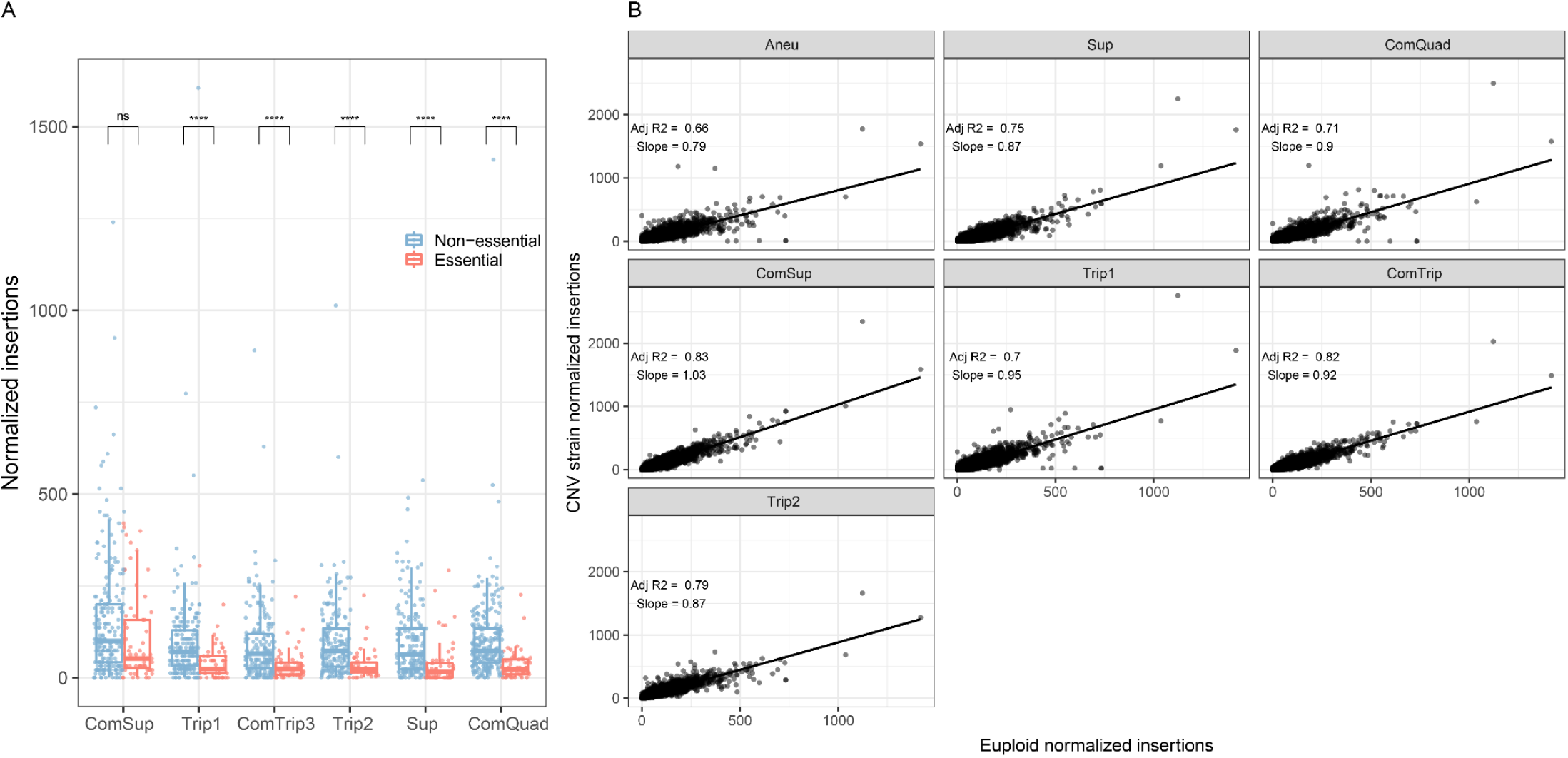
Transposon insertions in non-amplified genes. **A)** Boxplots of unique insertion sites per gene, with individual genes plotted as points, for essential (red) and non-essential (blue) genes (Winzeler et al. 1999). All genes on Chromosome XI that are not within the CNV boundaries are shown. P-values from Welch’s t-test are indicated by the following: ns: p > 0.01; ****: p < 0.0001. **B)** Linear regression was used to fit the normalized insertions per non-amplified gene in CNV strains (y-axis) to the mean number of normalized insertions per gene in the euploid replicates (x-axis), genome-wide. Adjusted p-values and slope from linear regression are annotated.

**Supplemental_Fig_S9.**
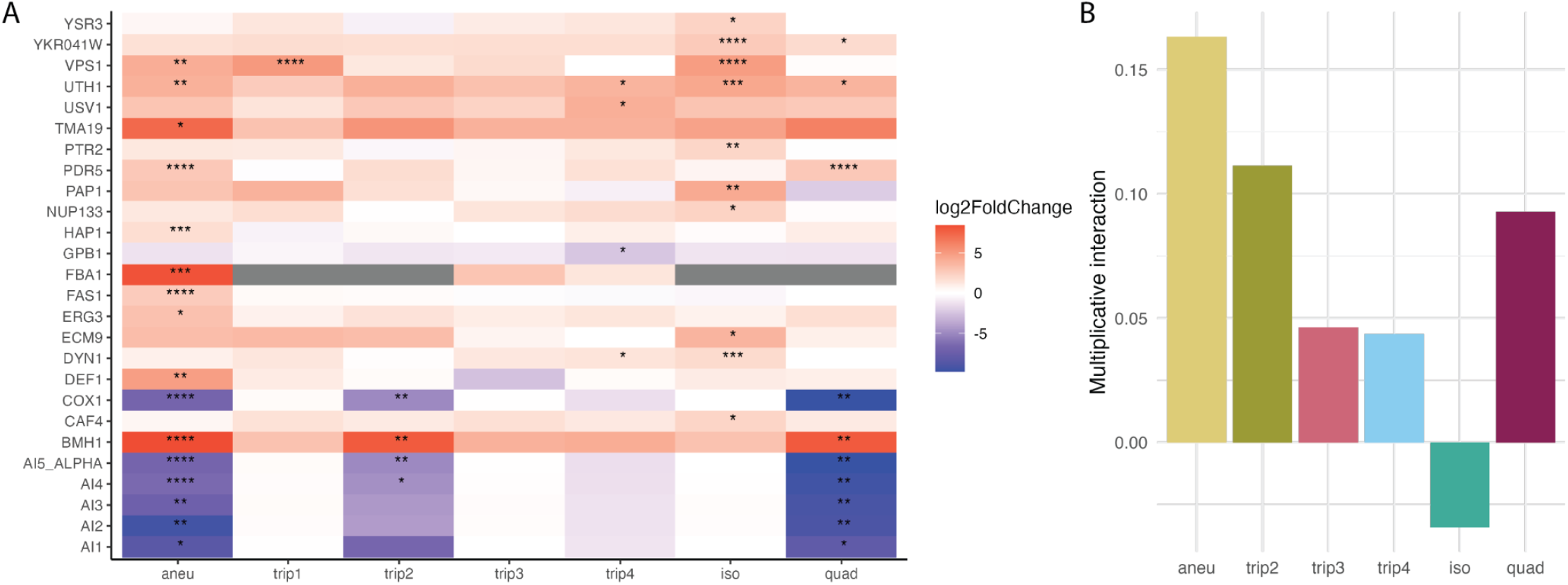
Genetic interactions of CNV strains. **A)** Genes that have significantly different insertions in CNV strains versus euploid. Genes which were significant for at least one CNV strain, from differential analysis comparing each CNV insertion profile to the euploid insertion profiles. Positive log2FoldChange values have more insertions in CNV strains than euploid strains, while negative log2FoldChange have fewer insertions in CNV strains than euploid strains. If a gene is amplified the copy number is annotated. P-values adjusted with the Benjamini and Hochberg method: *: p < 0.05; **: p < 0.01; ***: p < 0.001; ****: p < 0.0001. **B)** Multiplicative genetic interaction for CNV and *BMH1* double mutants. Calculated from growth rates shown in figure 3D.

**Supplemental_Fig_S10.**
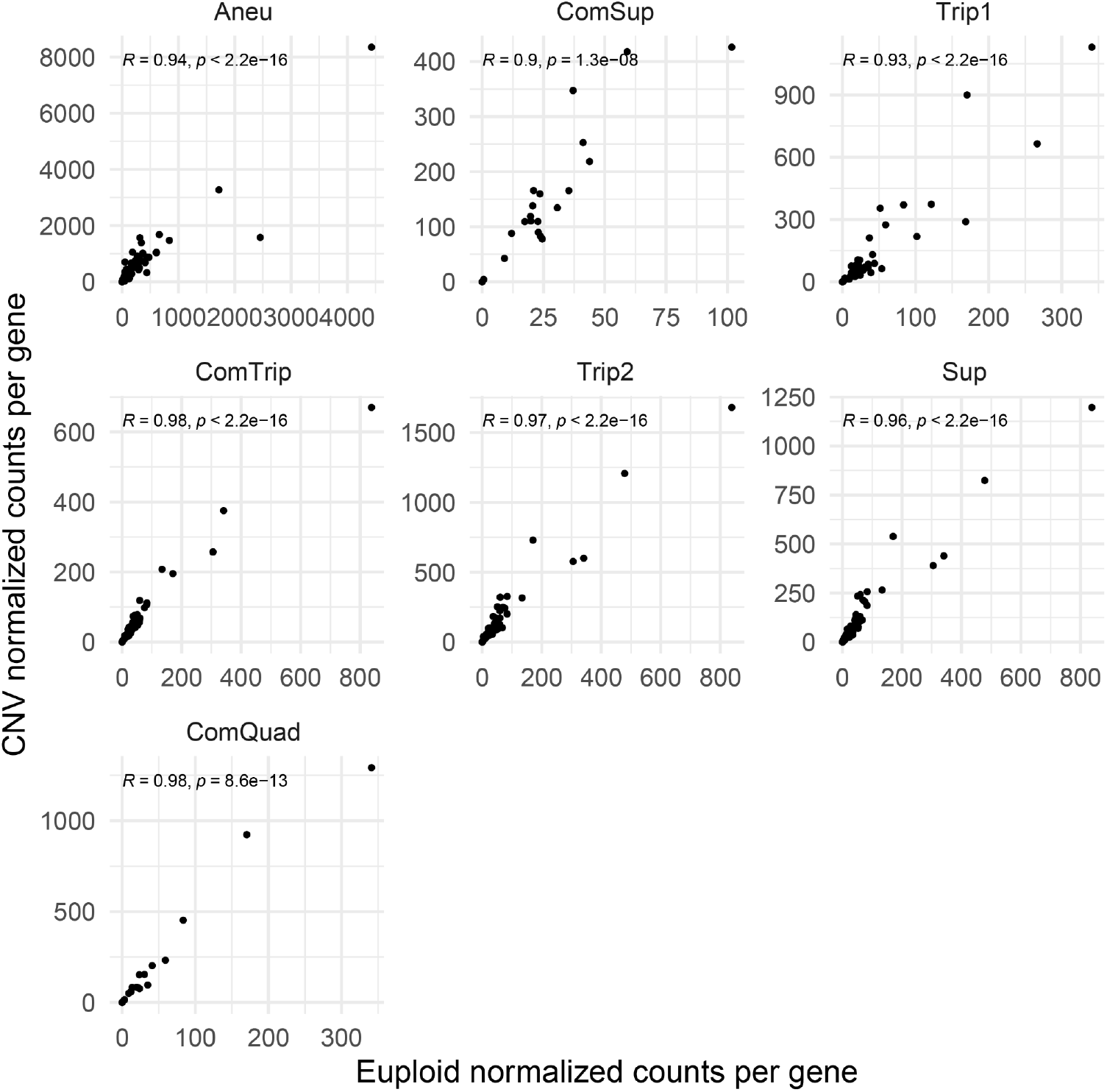
mRNA expression of amplified genes is highly correlated with euploid expression. For each CNV, the subset of genes within the CNV boundaries are shown. Pearson’s correlation coefficient and corresponding p-value are annotated.

**Supplemental_Fig_S11.**
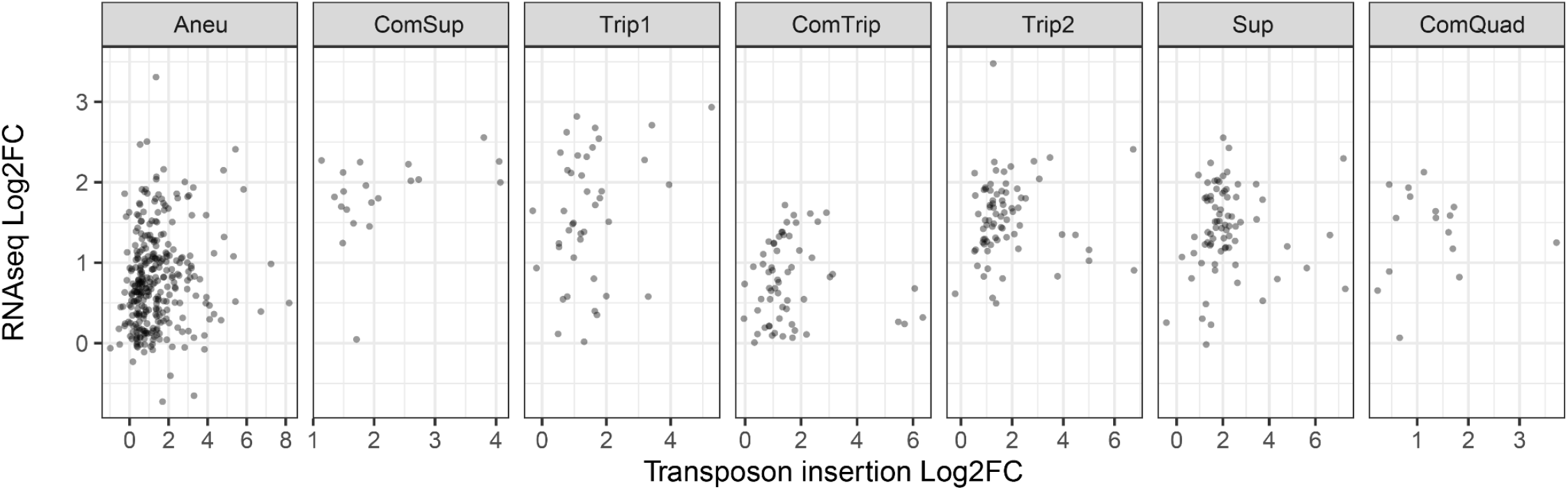
insertions not correlated with mRNA expression of amplified gene expression. For each CNV, the subset of genes within the CNV boundaries are shown. Pearson’s correlation coefficient and corresponding p-value are annotated.

**Supplemental_Fig_S12.**
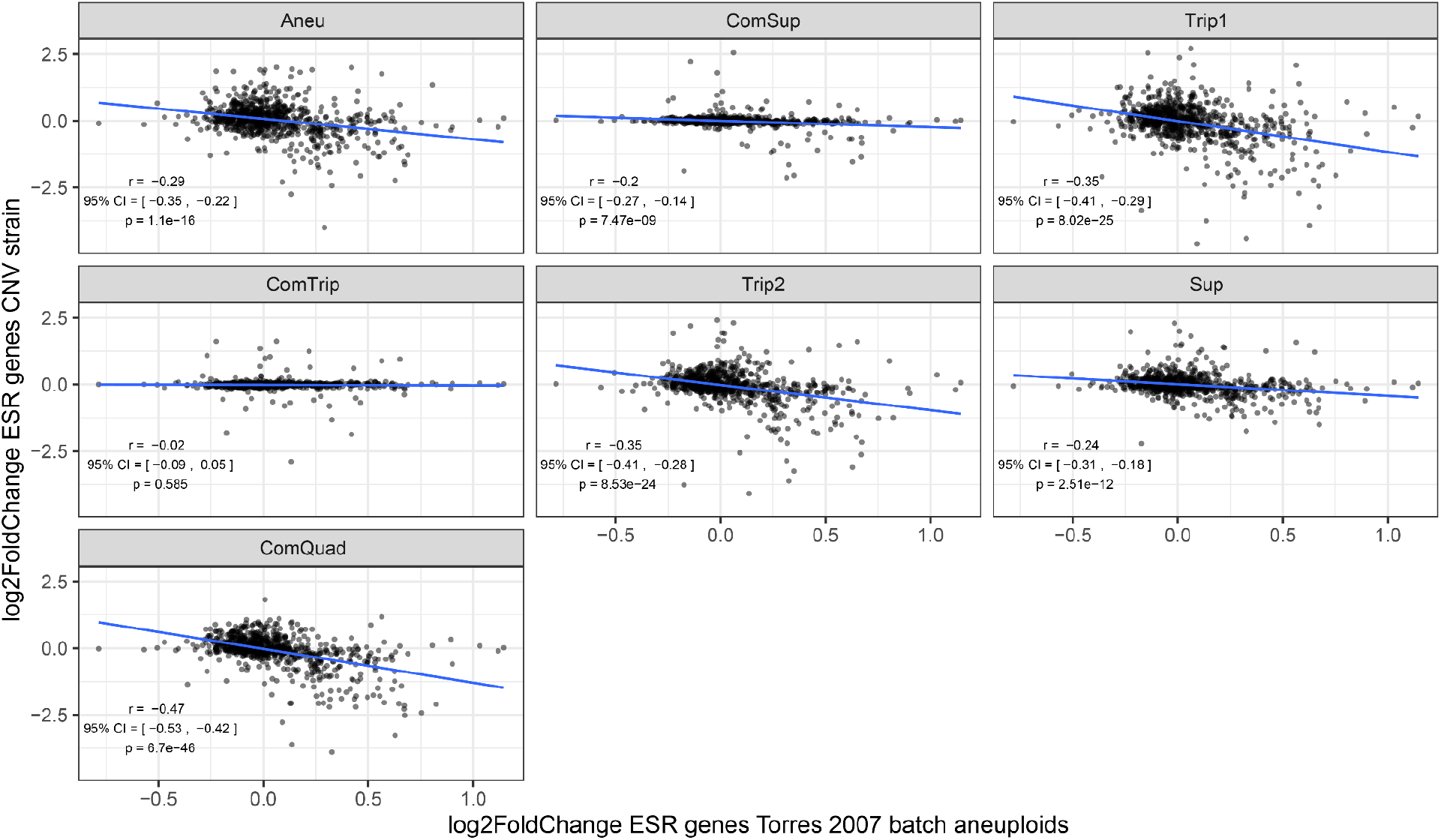
Pearson correlation between CNV strains and Torres 2007 aneuploids grown in batch culture for ESR genes. Log2 fold change in mRNA expression comparing CNV or aneuploid strain to euploid strain. The data from Torres is the mean for all aneuploid strains measured.

**Supplemental_Fig_S13.**
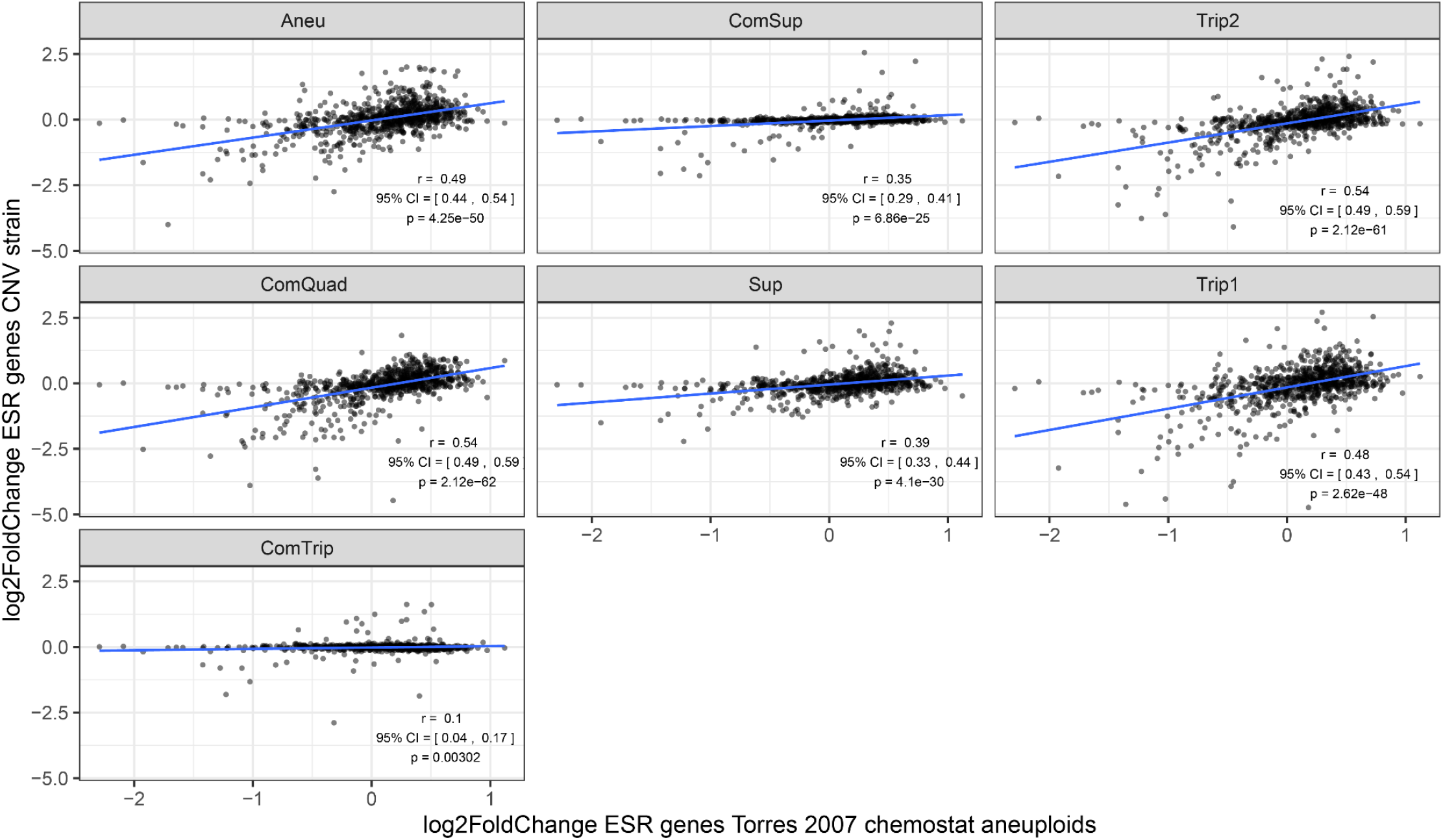
Pearson correlation between CNV strains and Torres 2007 aneuploids grown in chemostats for ESR genes. Log2 fold change in mRNA expression comparing CNV or aneuploid strain to euploid strain. The data from Torres is the mean for all aneuploid strains measured.

**Supplemental_Fig_S14.**
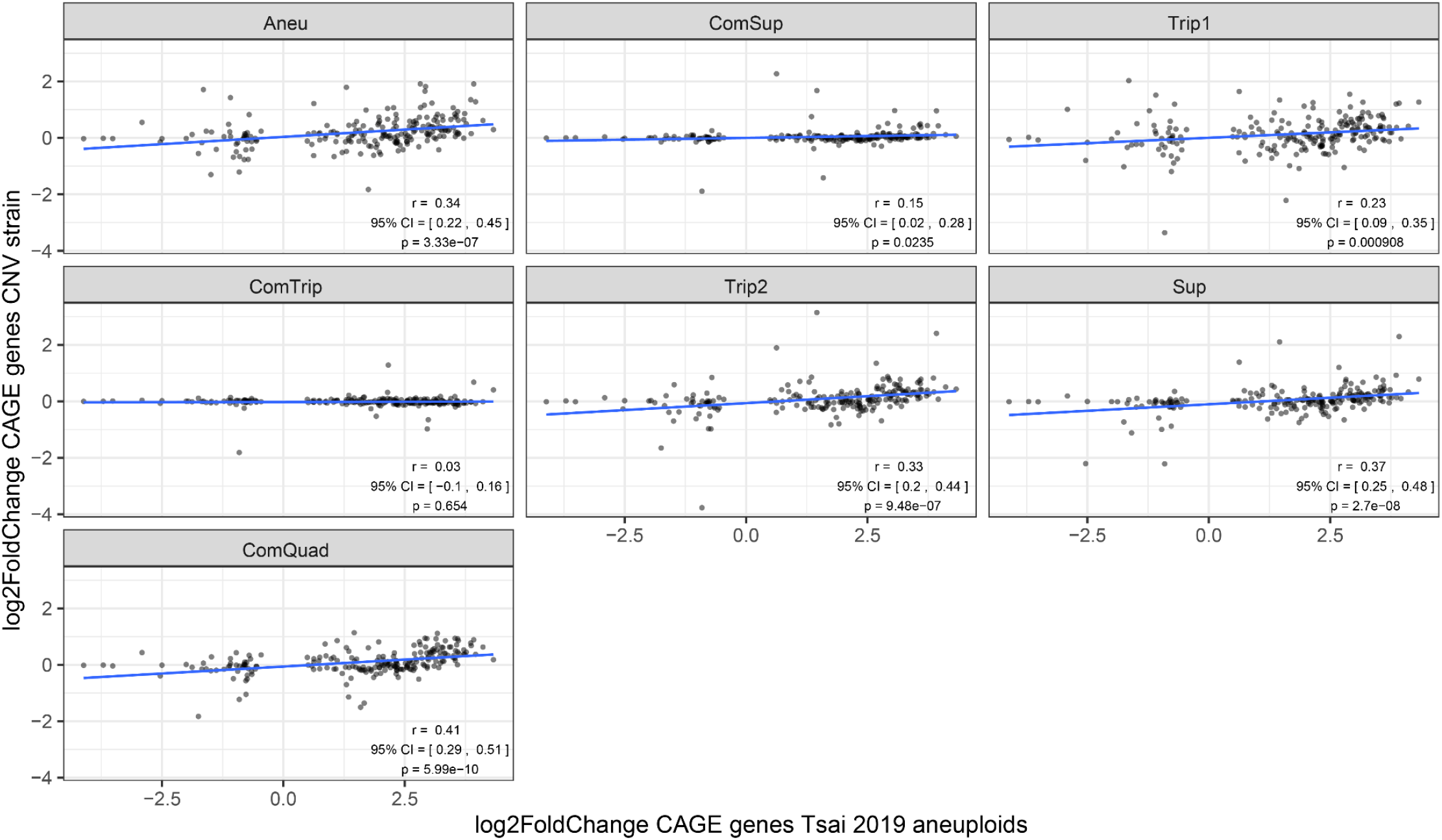
Pearson correlation between CNV strains and Tsai 2019 aneuploids for CAGE genes. Log2 fold change in mRNA expression comparing CNV or aneuploid strain to euploid strain.

**Supplemental_Fig_S15.**
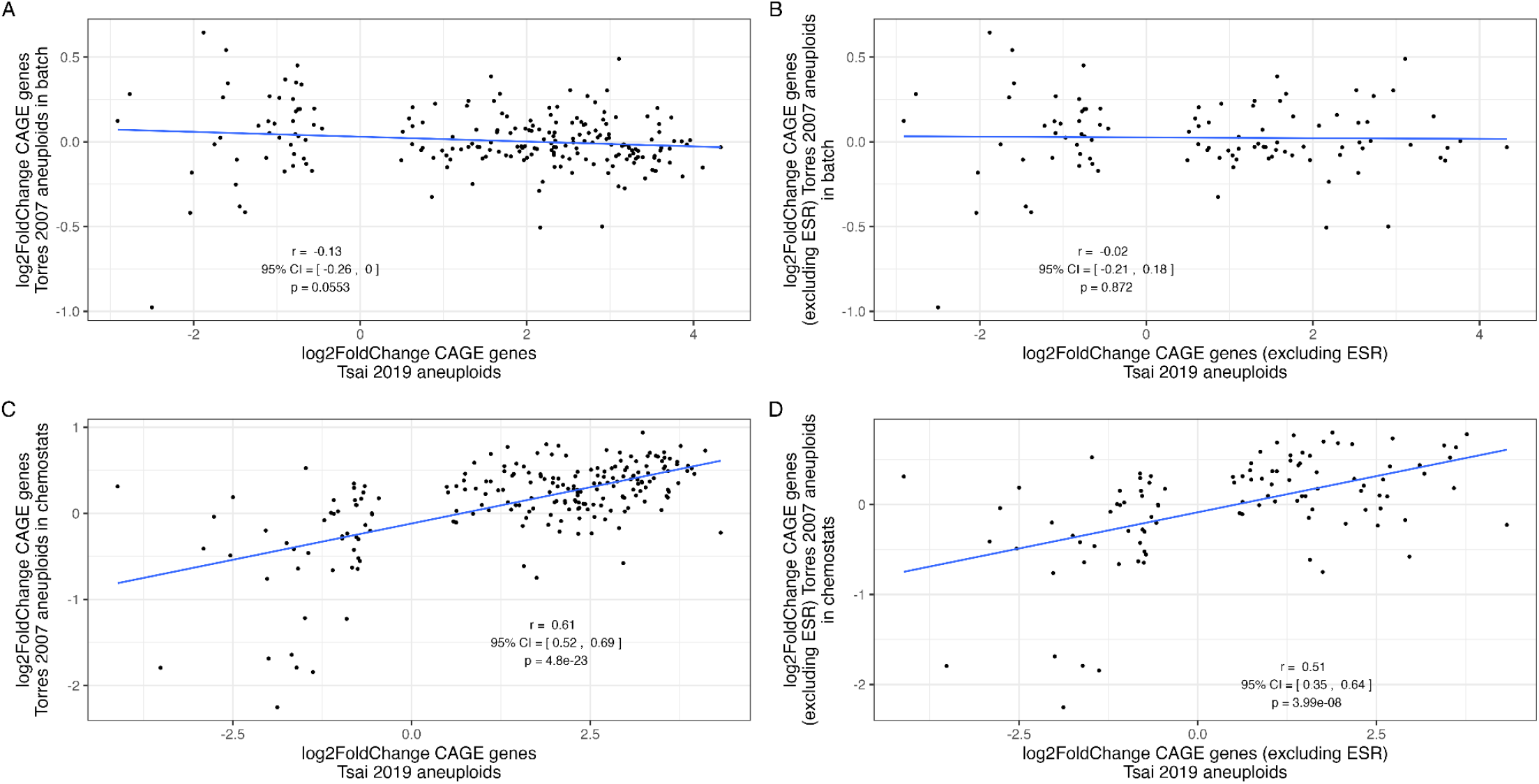
Pearson correlation between Torres 2007 aneuploids and Tsai 2019 aneuploids for CAGE genes. The data from Torres is the mean for all aneuploid strains measured.

**Supplemental_Fig_S16.**
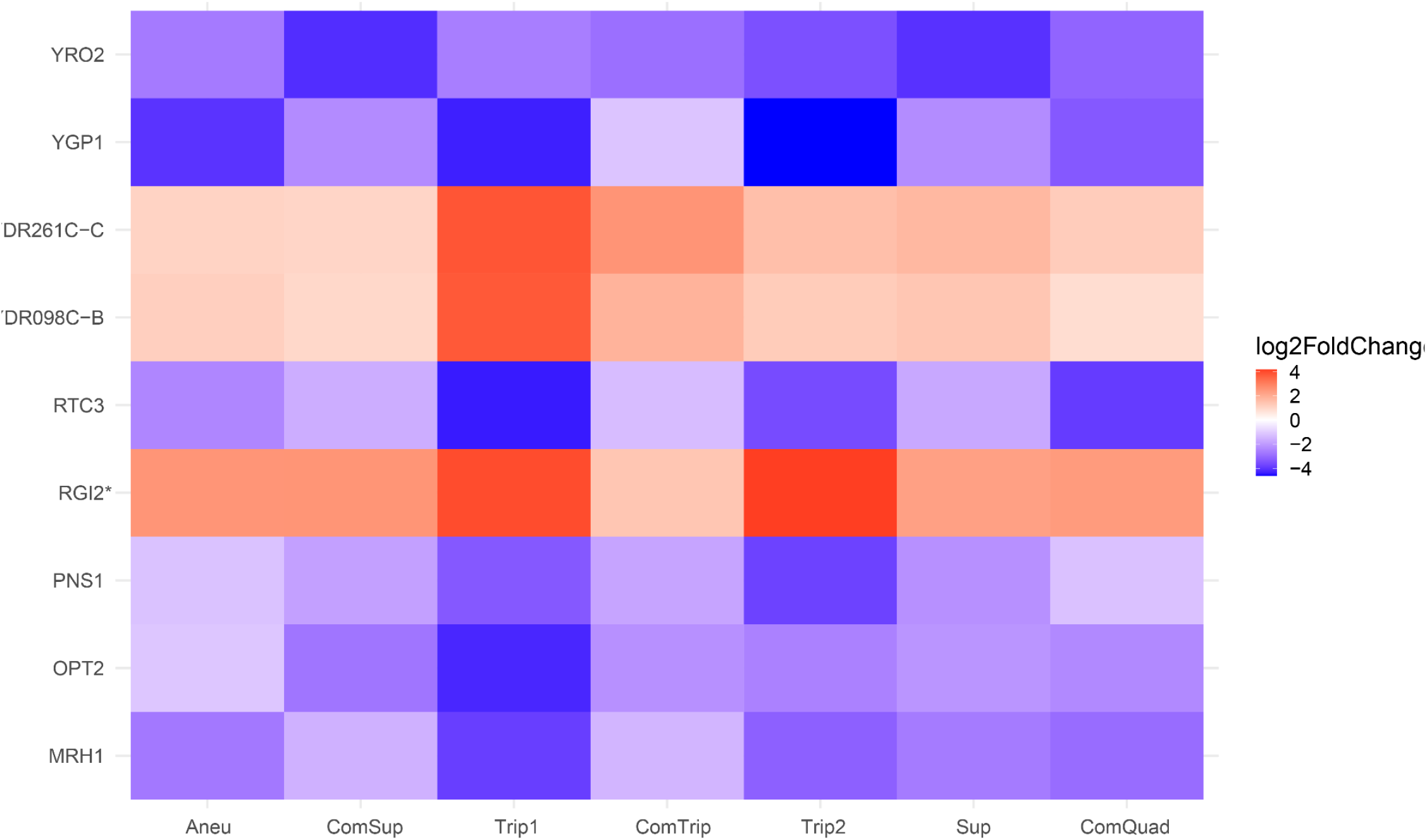
Genes with significantly different mRNA expression from the euploid in all strains that are not on chromosome XI. Genes with positive log2FoldChange have higher expression in the CNV strain than the euploid strain.

**Supplemental_Fig_S17.**
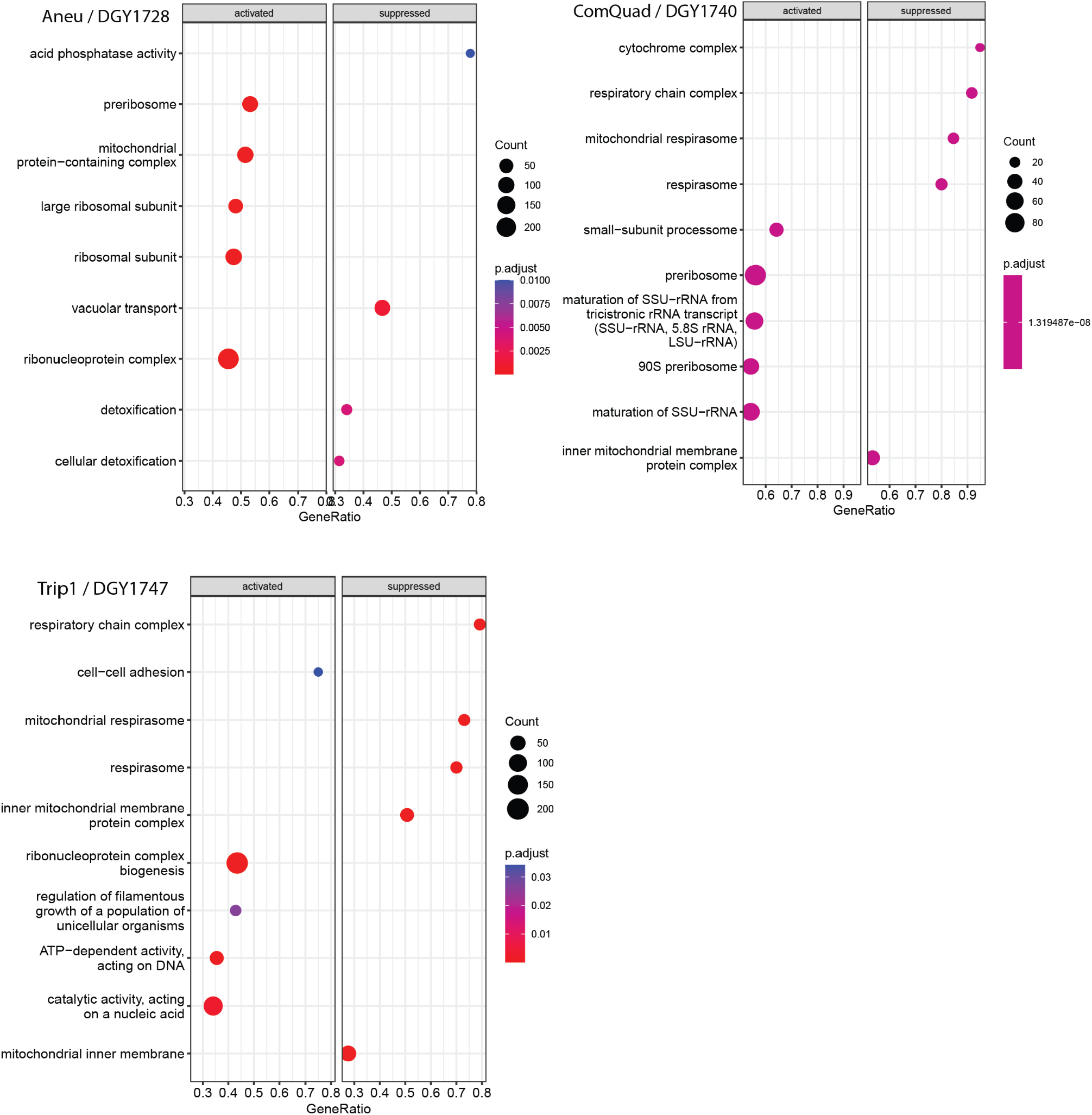
DESeq and GSEA differences between Trip1, ComQuad and Aneu. To better understand potential differences between the Trip1, ComQuad and Aneu strains we also performed DESeq to identify genes with significantly different mRNA abundances between these strains. We found that the most significant outlier in expression between these strains was INH1, a regulatory inhibitor of mitochondrial function, and SFT1, a INH1 paralog both associated with CCCP sensitivity (Ichikawa et al. 1990). Both of these are significantly higher in the Aneu strain (3.6 log2FC and 1.5 log2FC, respectively) and other CCCP resistant strains than in the CCCP sensitive strains. To help characterize how these strains may have distinct system level differences, we next identified genes with significantly different transcript abundances (DESeq2, adj.p-value <= 0.05) between each Aneu, Trip1, and ComQuad strain and the BMH1 insertion sensitive strains. A GSEA performed on these significantly different genes found that both ComQuad and Trip1 are enriched in suppressed ‘respiratory chain complex’, ‘respirasome / mitochondrial respirasome’ and ‘inner mitochondrial membrane protein complexes’. Intriguingly, Aneu alone is enriched in activating ‘mitochondrial protein-containing complexes’. This is suggestive of large scale differences in mitochondrial function particular to these strains.

## Supplemental Tables: All Supplementary tables are publicly available here

**https://github.com/pspealman/CNV_essentiality/tree/main/Supplementary_Tables/**

**Supplemental_Table_S1.** Strain characteristics. CNV type based on long read sequencing and genome assembly from (Spealman et al. 2022). More information about SNPs/indels including reference sequence and mutant sequence can be found in (Lauer et al. 2018) S10 Table https://doi.org/10.1371/journal.pbio.3000069.s027.

| Strain name | Strain name new | Gresham Lab Name | GAP1 hypothesized CNV type | Clone ID in Lauer et al. 2018 | Generation Isolated | SNPs/indels |
| --- | --- | --- | --- | --- | --- | --- |
| eu | Eu | DGY1657 | NA | NA | NA | NA |
| aneu | Aneu | DGY1728 | Aneuploid | gln_01_c1 | 150 | YNL284C-B missense variant; YPL232W (SSO1) disruptive inframe deletion |
| trip1 | ComSup | DGY1734 | Complex: supernumerary chromosome and ODIRA | gln_02_c3 | 250 | YHL002W (HSE1) missense variant; Chr XIV:96555 non-coding variant; Chr XIV:96603 non-coding variant |
| trip2 | Trip1 | DGY1747 | ODIRA | gln_08_c2 | 150 | YMR129W (POM152) missense variant; Chr V:431779 non-coding variant; Chr XII:915075 non-coding variant |
| trip3 | ComTrip | DGY1751 | Complex: ODIRA and homologous recombination | gln_09_c3 | 250 | YOL103W-A missense variant; YNR031C (SSK2) stop gained |
| trip4 | Trip2 | DGY1736 | ODIRA | gln_03_c2 | 250 | YJR152W (DAL5) stop lost & splice region variant & conservative inframe deletion; Chr V:55180 non-coding variant; Chr X:524178 non-coding variant; Chr X:745685 non-coding variant |
| iso | Sup | DGY1744 | Supernumerary chromosome | gln_07_c1 | 250 | YMR171C (EAR1) missense variant; YJL128C (PBS2) missense variant; Chr XV:594618 non-coding variant |
| quad | ComQuad | DGY1740 | Complex: ODIRA + transposon | gln_05_c1 | 150 | YOL077C (BRX1) missense variant; YNL338W frameshift_variant |

**Supplemental_Table_S2.** Change in genome size relative to ancestor.

| Name | Strain | CNV associated (bp) | rDNA locus (bp) | Total Change (bp) |
| --- | --- | --- | --- | --- |
| Aneu | DGY1728 | 666816 | -1149393 | -482577 |
| ComSup | DGY1734 | 372950 | -687962 | -315012 |
| Trip2 | DGY1736 | 321710 | 307378 | 629088 |
| ComQuad | DGY1740 | 79151 | -344556 | -265405 |
| Sup | DGY1744 | 396338 | -544994 | -148656 |
| Trip1 | DGY1747 | 191518 | -464468 | -272950 |
| ComTrip | DGY1751 | 168878 | -66842 | 102036 |

**Supplemental_Table_S3.** calculation of rDNA locus copy-number.

|  | DGY1657 | DGY1728 | DGY1734 | DGY1747 | DGY1751 | DGY1736 | DGY1744 | DGY1740 |
| --- | --- | --- | --- | --- | --- | --- | --- | --- |
| ETS1-1 | 83.5 | 36.3 | 61.0 | 65.5 | 82.0 | 94.4 | 65.0 | 73.5 |
| ETS1-2 | 69.2 | 32.8 | 44.1 | 50.8 | 61.2 | 70.8 | 50.9 | 57.5 |
| ETS2-1 | 65.9 | 19.9 | 33.3 | 42.8 | 45.3 | 59.5 | 44.9 | 39.0 |
| ETS2-2 | 84.5 | 32.8 | 62.4 | 68.5 | 82.9 | 95.5 | 66.0 | 71.2 |
| RDN18-1 | 80.9 | 32.0 | 58.9 | 64.7 | 78.6 | 89.7 | 61.8 | 70.8 |
| RDN18-2 | 71.7 | 30.2 | 51.3 | 57.2 | 68.2 | 76.9 | 54.6 | 62.9 |
| RDN25-1 | 68.7 | 26.6 | 47.0 | 53.2 | 61.6 | 70.7 | 50.5 | 55.2 |
| RDN25-2 | 82.4 | 29.9 | 61.0 | 66.1 | 80.9 | 89.4 | 63.8 | 67.5 |
| RDN37-1 | 73.0 | 28.5 | 51.4 | 58.5 | 70.7 | 79.0 | 55.1 | 63.0 |
| RDN37-2 | 78.3 | 30.0 | 55.6 | 63.1 | 77.0 | 86.0 | 60.4 | 66.2 |
| RDN5-1 | 84.3 | 33.8 | 63.1 | 69.7 | 83.8 | 93.9 | 65.8 | 73.3 |
| RDN5-2 | 65.7 | 17.9 | 28.9 | 40.7 | 42.3 | 56.0 | 43.1 | 35.7 |
| RDN58-1 | 76.8 | 29.6 | 53.3 | 62.1 | 74.3 | 85.2 | 59.0 | 67.2 |
| RDN58-2 | 77.7 | 29.2 | 55.0 | 63.3 | 76.3 | 86.5 | 60.5 | 67.4 |
| Median | 77.3 | 29.9 | 54.1 | 62.6 | 75.3 | 85.6 | 59.7 | 66.7 |
| Percentage | 1.000 | 0.387 | 0.701 | 0.810 | 0.975 | 1.108 | 0.773 | 0.863 |
| Size | 1351834 | 202441 | 663872 | 887366 | 1284992 | 1659212 | 806840 | 1007278 |
| Bp change | 0 | -1149393 | -687962 | -464468 | -66842 | 307378 | -544994 | -344556 |

**Supplemental_Table_S4.xlsx - Median Relative Depth of DNA per Gene**

**Supplemental_Table_S5.txt. - All strains Insertions per Gene, not normalized**

**Supplemental_Table_S6.txt. - All strains Insertions per Gene, normalized**

**Supplemental_Table_S7.txt - All strains Insertions per Gene, normalized**

**Supplemental_Table_S8.**
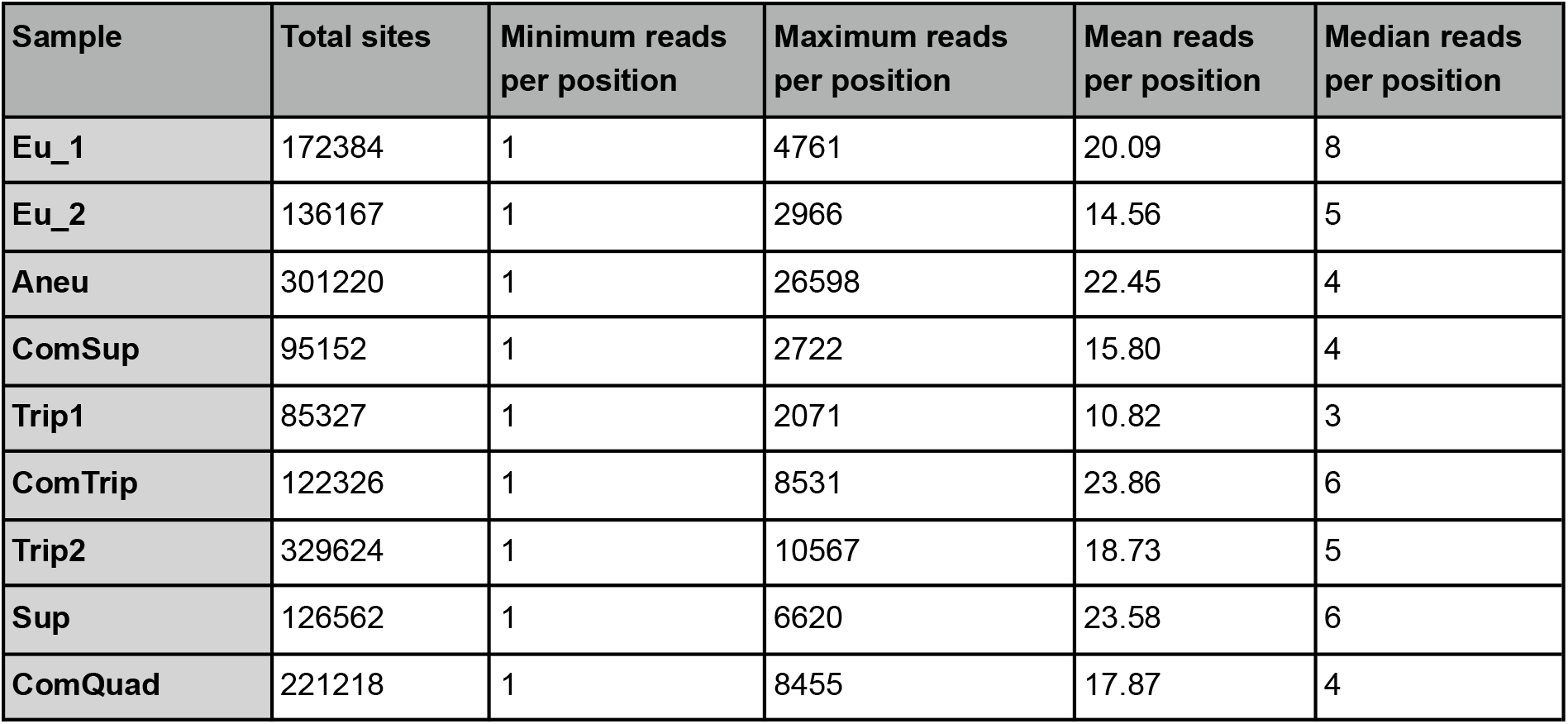
Hermes mutagenesis library characteristics for uniquely identified insertion sites.

**Supplemental_Table_S9.** Pearson correlation of insertions per gene for different sequencing runs.

|  | Euploid_1_nyc2 | Euploid_2_nyc1 | Euploid_2_nyc2 |
| --- | --- | --- | --- |
| Euploid_1_nyc1 | 0.979 | 0.918 | 0.903 |
| Euploid_1_nyc2 |  | 0.906 | 0.896 |
| Euploid_2_nyc1 |  |  | 0.966 |
|  | Aneu_bgi2 | Aneu_nyc1 |  |
| Aneu_bgi1 | 0.987 | 0.94 |  |
| Aneu_bgi2 |  | 0.94 |  |
|  | ComSup_nyc2 |  |  |
| ComSup_nyc1 | 0.957 |  |  |
|  | Trip1_nyc2 |  |  |
| Trip1_nyc1 | 0.904 |  |  |
|  | ComTrip_nyc2 |  |  |
| ComTrip_nyc1 | 0.966 |  |  |
|  | Trip2_bgi2 | Trip_nyc1 |  |
| Trip2_bgi1 | 0.974 | 0.928 |  |
| Trip2_bgi2 |  | 0.937 |  |
|  | Sup_nyc2 |  |  |
| Sup_nyc1 | 0.98 |  |  |
|  | ComQuad_bgi2 | ComQuad_nyc1 |  |
| ComQuad_bgi1 | 0.977 | 0.867 |  |
| ComQuad_bgi2 |  | 0.855 |  |

**Supplemental_Table_S10.txt - Summary of R-squared Outliers.** Tab-delimited table containing Normalized Tn abundances, copy-number, and standardized residuals for each gene in each strain relative to the euploid strain.

**Supplemental_Table_S11.txt - Calculation of number of genes exceeding R-squared significance threshold** Tab-delimited table containing the results of the DSG evaluation, namely, ‘cnv_hits’ is the times a CNV associated gene met the significance criteria (standardized residual > 2 and copy_number_corrected_log2FC > 1), ‘cnv_miss’ for when it failed those criteria. ‘Non_hits’ and ‘non_miss’ are the same test applied to non-CNV associated genes.

**Supplemental_Table_S12.csv - Genes with no insertions in euploid replicates.** These are genes that have no insertions in either replicate of the euploid strain 1657. If they were previously annotated as essential, they are labeled “yes” (Winzeler et al. 1999). Relative fitness for some of these genes on media with galactose was previously measured (column “Galactose”) (Costanzo et al. 2021), and are labeled as “low fitness galactose” if that relative fitness measure was less than one.

**Supplemental_Table_S13.xlsx - Gene set enrichment analysis of log2 fold change number of insertions per CNV strain compared to euploid replicates.** To simplify Figure 3B, we consolidated terms based on the overlap of the core enrichment between terms for each sample, the Keep column refers to if a term was kept for Figure 3B.

**Supplemental_Table_S14.xlsx - Results from differential analysis of number of insertions per CNV strain compared to euploid replicates.**

**Supplemental_Table_S15:** Pearson correlation of RNA abundance for different sequencing runs.

|  | Euploid_2 | Euploid_3 |
| --- | --- | --- |
| Euploid_1 | 0.989 | 0.995 |
| Euploid_2 | 1 | 0.984 |
|  | Aneu_2 | Aneu_3 |
| Aneu_1 | 0.996 | 0.955 |
| Aneu_2 | 1 | 0.956 |
|  | ComSup_2 | ComSup_3 |
| ComSup_1 | 0.981 | 0.937 |
| ComSup_2 | 1 | 0.974 |
|  | Trip_3 |  |
| Trip_1 | 0.991 |  |
|  | ComTrip_2 |  |
| ComTrip_1 | 0.999 |  |
|  | Trip2_2 | Trip_3 |
| Trip2_1 | 0.999 | 0.999 |
| Trip2_2 |  | 1 |
|  | Sup_2 | Sup_3 |
| Sup_1 | 0.998 | 1 |
| Sup_2 |  | 0.999 |
|  | ComQuad_2 | ComQuad_3 |
| ComQuad_1 | 0.237 | 1 |
| ComQuad_2 |  | 0.228 |

**Supplemental_Table_S16.txt - RNAseq read counts table for each strain.** Tab delimited results of BedTools coverage run of each sample, gene name corrected to Standard Name (SGD), only counting protein coding genes (Y*).

**Supplemental_Table_S17.txt - TPM normalized RNAseq abundances table for each strain.** The TPM normalized values of Supplemental_Table_S16.txt

**Supplemental_Table_S18.csv. Results from differential analysis of counts per gene from RNAseq for each CNV strain compared to euploid.**

**Supplemental_Table_S19.** STable 8 Mann-Whitney U test for Log2FoldChange of gene expression. This table shows the result of a Mann-Whitney U test comparing the log2 transformed TPM normalized RNA-seq abundances between the evolved CNV containing strains and the ancestral euploid strains. This test is conducted on the CNV associated genes and copy number normal genes. We find that the mean of CNV associated genes is significantly higher in the CNV strains than the euploid ancestor, with the CNV strains on average being 1.28 FC higher. Conversely non-CNV associated genes show no consistent FC across the CNV strains, with an average of 1.01 FC higher.

| CN_state | Strain | U_statistic | Evolved_Median | Ancestral_Median | Evo_over_Anc_ratio | p-value |
| --- | --- | --- | --- | --- | --- | --- |
| CNV | DGY1728 | 544154 | 5.960782 | 5.133703 | 1.161108 | 1.86E-16 |
| CNV | DGY1734 | 90467 | 5.921132 | 4.926497 | 1.201895 | 2.35E-10 |
| CNV | DGY1736 | 42994 | 6.737724 | 4.964693 | 1.357128 | 1.34E-26 |
| CNV | DGY1740 | 2498.5 | 6.808703 | 4.594912 | 1.481792 | 0.000249 |
| CNV | DGY1744 | 71967.5 | 6.239757 | 4.892394 | 1.2754 | 3.27E-18 |
| CNV | DGY1747 | 10934 | 6.419402 | 4.725971 | 1.358324 | 5.78E-08 |
| CNV | DGY1751 | 18748.5 | 5.819294 | 5.040278 | 1.154558 | 0.00013 |
| Non_CNV | DGY1728 | 1.36E+08 | 4.803955 | 4.917162 | 0.976977 | 1.85E-08 |
| Non_CNV | DGY1734 | 1.54E+08 | 5.040854 | 4.931223 | 1.022232 | 0.000134 |
| Non_CNV | DGY1736 | 1.64E+08 | 5.230348 | 4.930947 | 1.060719 | 1.72E-32 |
| Non_CNV | DGY1740 | 1.73E+08 | 5.355736 | 4.932343 | 1.08584 | 3.58E-70 |
| Non_CNV | DGY1744 | 1.55E+08 | 5.046671 | 4.932278 | 1.023193 | 6.25E-05 |
| Non_CNV | DGY1747 | 65623251 | 4.939313 | 5.06196 | 0.975771 | 1.73E-08 |
| Non_CNV | DGY1751 | 64253171 | 4.678993 | 4.905237 | 0.953877 | 1.17E-13 |

**Supplemental_Table_S20.txt -** Tab delimited file with copy number multiplied coverage counts from Supplemental_Table_S16.txt. Produced by PS_Supplemental_Script_repair_count_matrix.py.

**Supplemental_Table_S21.xlsx - DESeq2 results. Results from differential analysis of counts per gene from RNAseq for each CNV strain compared to euploid after copy number correction.**

**Supplemental_Table_S22.xlsx -** Table of FET analysis of DESeq2 results of CNV expression rates for both Observed and Expected values. Calculated using PS_Supplemental_Script_Calculate_rates_Obs_and_Exp.py. Observed data is the evolved strain compared to the ancestor with no copy number correction and we find significantly higher expression in CNV associated genes in each strain, compared to the ancestor, consistent with gene amplification models. Expected data is expression in the CNV strain compared to copy number corrected ancestor expression. We do not find a significant difference in expression of CNV associated genes from what is expected given their copy number, suggesting there is no dosage compensation specific to CNVs.

**Supplemental_Table_S23.csv - Gene set enrichment analysis of log2 fold change counts per gene from RNAseq for each CNV strain compared to euploid.**

**Supplemental_Table_S24.csv - Results of hypergeometric test for over-representation of GO terms in clustered, differentially expressed genes.**

**Supplemental_Table_S25.csv - STable 10 Results of hypergeometric test for over-representation of GO terms in clustered, differentially expressed genes excluding genes on chromosome XI.**

**Supplemental_Table_S26.**
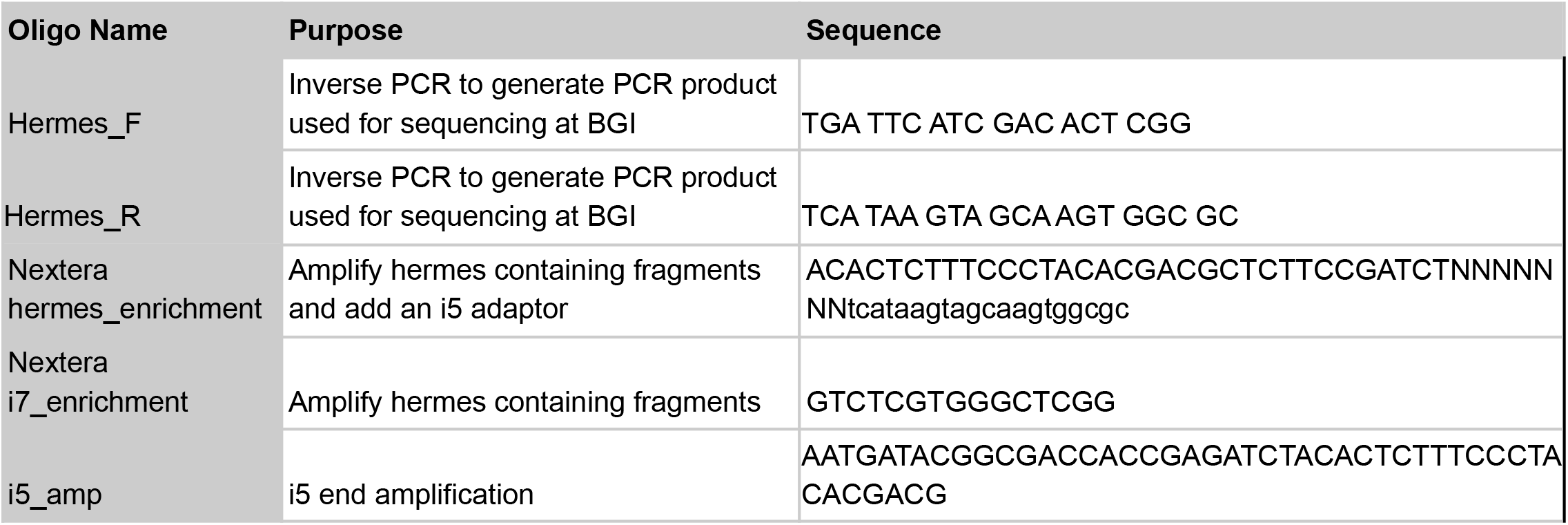
Oligos and Primers used in this work:

## Supplemental Files

**All Supplemental files are publicly available here:**

https://github.com/pspealman/CNV_essentiality/tree/main/Supplemental_Files/

**Supplemental_File_S1.zip - Modified reference genome GCF_000146045.2_R64_genomic_GAP1.gff**

Modified GCF_000146045.2_R64_genomic.gff file with the Gresham GFP Reporter added to the appropriate coordinates on ChrXI.

**Supplemental_File_S2.html - Description of source supplementary data present in Supplemental_File_S3.zip**

**Supplemental_File_S3.zip - Collection of custom source code used in the analysis and generation of data.** Source code and supplementary data as described in Supplemental_File_S2.html.

## Acknowledgements

We thank members of the Gresham, Schacherer and Vogel Lab. This work was made possible by grants from the NIH to P.S. (F32-GM131573) and D.G. (R01-GM134066, R01-GM107466) and from the NSF to G.A. (DGE1342536) and DG (MCB1818234). J.S. is supported by the European Research Council (ERC) Consolidator grant (772505). J.S. is a fellow of the University of Strasbourg Institute for Advanced Study (USIAS) and a member of the Institut Universitaire de France.

## Notes

### Competing Interest Statement

The authors have declared no competing interest.

