## Supplemental File 2 for "Copy number variants alter local and global mutational tolerance"

Description of code base for CNV\_essentiality paper


### Description of code base for CNV\_essentiality paper

###### Pieter Spealman

#### 2022-12-15

###### Generate Insertion position data from Tn FASTQ

```
#Scripts that were used to generate insertion position data are all in 

onHPC/onHPC_fastq_to_insertions.Rmd
```

###### Script used to QC, align, and count RNAseq data

```
#Scripts that were used to QC, align, and count RNAseq data  

onHPC/run_windchime_star.sh
```

###### Script used to QC, align, and count RNAseq data

```
#analysis/generate figures for the transposon insertion data

analysis/functions.R
analysis/growth_rate_summary.Rmd
analysis/hermes_analysis.Rmd
```
