## Supplemental File 3 for "Copy number variants alter local and global mutational tolerance": compare_rna_hermes.Rmd

````
---
title: "R Notebook"
output: html_notebook
---

This is an [R Markdown](http://rmarkdown.rstudio.com) Notebook. When you execute code within the notebook, the results appear beneath the code. 

Try executing this chunk by clicking the *Run* button within the chunk or by placing your cursor inside it and pressing *Cmd+Shift+Enter*. 

```{r}
con_insert_profiles = read_csv(paste0(data_dir,"/conservative_insertion_profiles.csv"))

counts_df = read_tsv(paste0(data_dir, "/coverage_table_rnaseq.txt")) %>% mutate(gene = `#gene`) %>%
  filter(!is.na(gene))
# convert "genex" to systematic id
for(i in 1:nrow(counts_df)) {
  y = gff$X9[str_detect(gff$X9, paste0("ID=",counts_df$gene[i]))] %>% str_split(";")
  z = str_remove(y[[1]][str_detect(y[[1]],"locus_tag")],"locus_tag=")
  counts_df$gene[i] = z
}
counts_df = counts_df %>% select(-`#gene`)

counts_rnaseq = counts_df %>%
  pivot_longer(cols=-gene, names_to="names", values_to = "rnaseq_counts") %>%
  mutate(sample = str_sub(names, 4, -3)) %>%
  group_by(sample, gene) %>%
  summarise(mean_rnaseq_counts = mean(rnaseq_counts)) %>% ungroup() 

all_data = counts_rnaseq %>% 
  right_join(con_insert_profiles %>% filter(type=="cds") %>% filter(sample != "1657_2") %>% mutate(sample = if_else(sample == "1657_1", "1657", sample)))

# interactive?
library(plotly)
p=ggplot(all_data, aes(mean_rnaseq_counts, n_insertions, text = gene)) +
  scale_color_manual(values = "black") +
  geom_point() +
  facet_wrap(~sample) #+
  scale_y_log10() +
  scale_x_log10()
ggplotly(p)
```
````
