## Supplemental File 3 for "Copy number variants alter local and global mutational tolerance": Description_of_Code.rmd

````
---
title: "Description of code base for CNV_essentiality paper"
author: "Pieter Spealman"
date: "`r Sys.Date()`"
output: html_document
---
##### Generate Insertion position data from Tn FASTQ
```{}
#Scripts that were used to generate insertion position data are all in 

onHPC/onHPC_fastq_to_insertions.Rmd
```

##### Script used to QC, align, and count RNAseq data

```{}
#Scripts that were used to QC, align, and count RNAseq data  

onHPC/run_windchime_star.sh
```

##### Script used to QC, align, and count RNAseq data

```{}
#analysis/generate figures for the transposon insertion data

analysis/functions.R
analysis/growth_rate_summary.Rmd
analysis/hermes_analysis.Rmd

```

##### Script used to generate some of the RNAseq analysis and plots

```{}
#analysis/generate figures for the RNAseq data

rnaseq_r/rnaseq_analysis.Rmd

```

##### Scripts used in Tnseq copy_number_corrected re-analysis 

```{}
#relabel genes and perform unique count normalization

tnseq_cnc/tiny_insert_counter.py

#Make Fig2B and Insert Heatmap supplemental file

tnseq_cnc/Make_Insert_Heatmap.py

#Calculate R-squared linear regression and significant residual outliers 

tnseq_cnc/Make_Insert_R-squared.r

#Identify CNV-associated genes that are significantly higher in all the strains they are amplified in.

tnseq_cnc/Make_Supplemental_Fig2C_threshold.py

```

##### Scripts used in RNAseq copy_number_corrected re-analysis 

```{}
#relabel genes and set expected (Exp) expression values using the Observed (Obs) expression, namely the Obs RNA abundance of the ancestor mulitplied by the observed gene copy number

rnaseq_cnc/repair_count_matrix.py

#DESeq2 code using the observed (Obs) expression values

rnaseq_cnc/DESeq_difObs_v1.0.R

#DESeq2 code using the expected (Exp) expression values

rnaseq_cnc/DESeq_difExp_v1.0.R

#Calculate rates of significant gene expression differences in the Obs and Exp groups

rnaseq_cnc/Calculate_rates_Obs_and_Exp.py

#Make figure 4 subfigures:

rnaseq_cnc/Make_Fig4.py
rnaseq_cnc/Make_Fig4v2_DESeq.py
rnaseq_cnc/Make_Fig4C.py

#Calculate if CNV associated genes have a higher RNA abundance than their non-CNV ancestors (tpm normalized). Mann-Whitney U test for Log2FoldChange of gene expression.

rnaseq_cnc/Make_Sup_Compare_CNV_to_Non.py

```
````
