## Supplemental File 3 for "Copy number variants alter local and global mutational tolerance": Make_Insert_R-squared.r

```
global_normalized_insertionPerGene <- read.delim("C:/Gresham/tiny_projects/Project_Grace/insertions/global_both_eu_normalized_insertionPerGene.txt")
relative_depth_DNA_corrected_v3 <- read.delim("C:/Gresham/tiny_projects/Project_Grace/relative_depth_DNA_corrected_v3.txt")

###
##pdf("C:/Gresham/tiny_projects/Project_Grace/supplemental_figures/Supplemental_Fig2C_all_combined_DGY1657.pdf")
combined <- merge(global_normalized_insertionPerGene, relative_depth_DNA_corrected_v3, by.x = "X", by.y = "Gene")
combined <- na.omit(combined)

cols <- rep(rgb(0,0,0,0.01),length(combined$X)) # initialize cols to black
cols[combined$DGY1657==2] <- rgb(1,0,0,0.5)
cols[combined$DGY1657==3] <- rgb(0,0,1,0.5)
cols[combined$DGY1657==4] <- rgb(0,1,1,0.5)

pchs <- rep(16, length(combined$X)) # initialize cols filled circle
pchs[combined$DGY1657==0] <- 3
pchs[combined$DGY1657==3] <- 17
pchs[combined$DGY1657==4] <- 18

ismax <- max(combined$X1657_1, combined$X1657_2)

# ismax <- max(combined$X1657, combined$X1728, combined$X1734, combined$X1736,
#              combined$X1740, combined$X1744, combined$X1747, combined$X1751,
#              na.rm =TRUE) 

plot(combined$X1657_1, combined$X1657_2, 
     col = cols, pch = pchs, xlim=c(0, ismax), ylim = c(0,ismax))

fit<-lm(combined$X1657_2~combined$X1657_1)
abline(fit)
summary(fit)

combined$standardized_residuals<-rstandard(fit)
res_cutoff<-2*sd(combined$standardized_residuals)+mean(combined$standardized_residuals)
DGY_hot<- subset(combined, abs(combined$standardized_residuals) > 2)

write.table(combined,
            'C:/Gresham/tiny_projects/Project_Grace/supplemental_figures/Supplemental_Fig2C_Outliers_CNV_DGY1657.txt', 
            sep='\t', row.names = FALSE, quote = FALSE)

points(DGY_hot$X1657_1, DGY_hot$X1657_2, col=rgb(1,0,0,0.5), pch = 0)

#DGY_ <- subset(combined, combined$DGY1657!=1)
#fit<-lm((DGY_$X1657_2)~((DGY_$X1657_1)))
#abline(fit, col='red')
#summary(fit)

#dev.off()

x<-subset(DGY_hot, DGY_hot$DGY1657!=1)
y<-subset(combined, combined$DGY1657!=1)
m<-subset(DGY_hot, DGY_hot$DGY1657==1)
n<-subset(combined, combined$DGY1657==1)


DGY_ <- subset(combined, combined$DGY1657!=1)
cols <- rep(rgb(0,0,0,0.2),length(DGY_)) # initialize cols to black
cols[DGY_$DGY1657==2] <- rgb(0,0,0,0.2)
cols[DGY_$DGY1657==3] <- rgb(0,1,0,0.2)
cols[DGY_$DGY1657==4] <- rgb(0,0,1,0.2)

pchs <- rep(16, length(DGY_)) # initialize cols filled circle
pchs[DGY_$DGY1657==0] <- 3
pchs[DGY_$DGY1657==3] <- 2
pchs[DGY_$DGY1657==4] <- 5

points((DGY_$X1657_1), (DGY_$X1657_2), 
       col = cols, xlim=c(0, ismax), ylim = c(0,ismax), pch = pchs)

plot((DGY_$X1657_1), (DGY_$X1657_2), 
     col = cols, xlim=c(0, ismax), ylim = c(0,ismax), pch = pchs)
fit<-lm((DGY_$X1657_2)~((DGY_$X1657_1)))
abline(fit, col='red')
summary(fit)

DGY_$standardized_residuals<-rstandard(fit)
res_cutoff<-2*sd(DGY_$standardized_residuals)+mean(DGY_$standardized_residuals)
DGY_hot<- subset(DGY_, abs(DGY_$standardized_residuals) > 2)

points(DGY_hot$X1657_1, DGY_hot$X1657_2, col=rgb(1,0,0,0.5), pch = 0)

###
global_normalized_insertionPerGene <- read.delim("C:/Gresham/tiny_projects/Project_Grace/insertions/global_normalized_insertionPerGene_noDGY.txt")
relative_depth_DNA_corrected_v3 <- read.delim("C:/Gresham/tiny_projects/Project_Grace/relative_depth_DNA_corrected_v3.txt")

###
#pdf("C:/Gresham/tiny_projects/Project_Grace/supplemental_figures/Supplemental_Fig2C_all_combined_DGY1728.pdf")
combined <- merge(global_normalized_insertionPerGene, relative_depth_DNA_corrected_v3, by.x = "X", by.y = "Gene")
combined <- na.omit(combined)

cols <- rep(rgb(0,0,0,0.01),length(combined$X)) # initialize cols to black
cols[combined$DGY1728==2] <- rgb(1,0,0,0.5)
cols[combined$DGY1728==3] <- rgb(0,0,1,0.5)
cols[combined$DGY1728==4] <- rgb(0,1,1,0.5)

pchs <- rep(16, length(combined$X)) # initialize cols filled circle
pchs[combined$DGY1728==0] <- 3
pchs[combined$DGY1728==3] <- 17
pchs[combined$DGY1728==4] <- 18

ismax <- max(combined$X1657, combined$X1728)

# ismax <- max(combined$X1657, combined$X1728, combined$X1734, combined$X1736,
#              combined$X1740, combined$X1744, combined$X1747, combined$X1751,
#              na.rm =TRUE) 

plot(combined$X1657, combined$X1728, 
     col = cols, pch = pchs)

fit<-lm(combined$X1728~combined$X1657)
abline(fit)
summary(fit)

combined$standardized_residuals<-rstandard(fit)
res_cutoff<-2*sd(combined$standardized_residuals)+mean(combined$standardized_residuals)
DGY_hot<- subset(combined, abs(combined$standardized_residuals) > 2)

write.table(DGY_hot,
            'C:/Gresham/tiny_projects/Project_Grace/supplemental_figures/Supplemental_Fig2C_SigOutliers_DGY1728.txt', 
            sep='\t', row.names = FALSE, quote = FALSE)

write.table(combined,
            'C:/Gresham/tiny_projects/Project_Grace/supplemental_figures/Supplemental_Fig2C_Outliers_CNV_DGY1728.txt', 
            sep='\t', row.names = FALSE, quote = FALSE)
            

points(DGY_hot$X1657, DGY_hot$X1728, col=rgb(1,0,0,0.5), pch = 0)

DGY_ <- subset(combined, combined$DGY1728!=1)
fit<-lm((DGY_$X1728)~((DGY_$X1657)))
abline(fit, col='red')
summary(fit)

dev.off()

x<-subset(DGY_hot, DGY_hot$DGY1728!=1)
y<-subset(combined, combined$DGY1728!=1)
m<-subset(DGY_hot, DGY_hot$DGY1728==1)
n<-subset(combined, combined$DGY1728==1)


DGY_ <- subset(combined, combined$DGY1728!=1)
cols <- rep(rgb(0,0,0,0.2),length(DGY_)) # initialize cols to black
cols[DGY_$DGY1728==2] <- rgb(0,0,0,0.2)
cols[DGY_$DGY1728==3] <- rgb(0,1,0,0.2)
cols[DGY_$DGY1728==4] <- rgb(0,0,1,0.2)

pchs <- rep(16, length(DGY_)) # initialize cols filled circle
pchs[DGY_$DGY1728==0] <- 3
pchs[DGY_$DGY1728==3] <- 2
pchs[DGY_$DGY1728==4] <- 5

points((DGY_$X1657), (DGY_$X1728), 
       col = cols, xlim=c(0, ismax), ylim = c(0,ismax), pch = pchs)

plot((DGY_$X1657), (DGY_$X1728), 
     col = cols, xlim=c(0, ismax), ylim = c(0,ismax), pch = pchs)
fit<-lm((DGY_$X1728)~((DGY_$X1657)))
abline(fit, col='red')
summary(fit)

DGY_$standardized_residuals<-rstandard(fit)
res_cutoff<-2*sd(DGY_$standardized_residuals)+mean(DGY_$standardized_residuals)
DGY_hot<- subset(DGY_, abs(DGY_$standardized_residuals) > 2)

write.table(DGY_hot,
            'C:/Gresham/tiny_projects/Project_Grace/supplemental_figures/Supplemental_Fig2C_SigOutliers_CNV_DGY1728.txt', 
            sep='\t', row.names = FALSE, quote = FALSE)

write.table(DGY_,
            'C:/Gresham/tiny_projects/Project_Grace/supplemental_figures/Supplemental_Fig2C_Outliers_CNV.Only_DGY1728.txt', 
            sep='\t', row.names = FALSE, quote = FALSE)


points(DGY_hot$X1657, DGY_hot$X1728, col=rgb(1,0,0,0.5), pch = 0)

####
#pdf("C:/Gresham/tiny_projects/Project_Grace/supplemental_figures/Supplemental_Fig2C_all_combined_DGY1740.pdf")
combined <- merge(global_normalized_insertionPerGene, relative_depth_DNA_corrected_v3, by.x = "X", by.y = "Gene")
combined <- na.omit(combined)

DGY_ <- subset(combined, combined$DGY1740!=1)
cols <- rep(rgb(0,0,0,0.01),length(combined$X)) # initialize cols to black
cols[combined$DGY1740==2] <- rgb(1,0,0,0.5)
cols[combined$DGY1740==3] <- rgb(0,0,1,0.5)
cols[combined$DGY1740==4] <- rgb(0,1,1,0.5)

pchs <- rep(16, length(combined$X)) # initialize cols filled circle
pchs[combined$DGY1740==0] <- 3
pchs[combined$DGY1740==3] <- 17
pchs[combined$DGY1740==4] <- 18

ismax <- max(combined$X1657, combined$X1740)

# ismax <- max(combined$X1657, combined$X1728, combined$X1734, combined$X1736,
#              combined$X1740, combined$X1744, combined$X1747, combined$X1751,
#              na.rm =TRUE) 

plot(combined$X1657, combined$X1740, 
     col = cols, pch = pchs, xlim=c(0, ismax), ylim = c(0,ismax))

fit<-lm(combined$X1740~combined$X1657)
abline(fit)
summary(fit)

combined$standardized_residuals<-rstandard(fit)
res_cutoff<-2*sd(combined$standardized_residuals)+mean(combined$standardized_residuals)
DGY_hot<- subset(combined, abs(combined$standardized_residuals) > 2)

write.table(DGY_hot,
            'C:/Gresham/tiny_projects/Project_Grace/supplemental_figures/Supplemental_Fig2C_SigOutliers_DGY1740.txt', 
            sep='\t', row.names = FALSE, quote = FALSE)

write.table(combined,
            'C:/Gresham/tiny_projects/Project_Grace/supplemental_figures/Supplemental_Fig2C_Outliers_CNV_DGY140.txt', 
            sep='\t', row.names = FALSE, quote = FALSE)


points(DGY_hot$X1657, DGY_hot$X1740, col=rgb(1,0,0,0.5), pch = 0)

fit<-lm((DGY_$X1740)~((DGY_$X1657)))
abline(fit, col='red')
summary(fit)

dev.off()

x<-subset(DGY_hot, DGY_hot$DGY1740!=1)
y<-subset(combined, combined$DGY1740!=1)
m<-subset(DGY_hot, DGY_hot$DGY1740==1)
n<-subset(combined, combined$DGY1740==1)


DGY_ <- subset(combined, combined$DGY1740!=1)
cols <- rep(rgb(0,0,0,0.2),length(DGY_)) # initialize cols to black
cols[DGY_$DGY1740==2] <- rgb(0,0,0,0.2)
cols[DGY_$DGY1740==3] <- rgb(0,1,0,0.2)
cols[DGY_$DGY1740==4] <- rgb(0,0,1,0.2)

pchs <- rep(16, length(DGY_)) # initialize cols filled circle
pchs[DGY_$DGY1740==0] <- 3
pchs[DGY_$DGY1740==3] <- 2
pchs[DGY_$DGY1740==4] <- 5

points((DGY_$X1657), (DGY_$X1740), 
     col = cols, xlim=c(0, ismax), ylim = c(0,ismax), pch = pchs)

plot((DGY_$X1657), (DGY_$X1740), 
     col = cols, xlim=c(0, ismax), ylim = c(0,ismax), pch = pchs)
fit<-lm((DGY_$X1740)~((DGY_$X1657)))
abline(fit, col='red')
summary(fit)

DGY_$standardized_residuals<-rstandard(fit)
DGY_hot<- subset(DGY_, abs(DGY_$standardized_residuals) > 2)

write.table(DGY_hot,
            'C:/Gresham/tiny_projects/Project_Grace/supplemental_figures/Supplemental_Fig2C_SigOutliers_CNV_DGY1740.txt', 
            sep='\t', row.names = FALSE, quote = FALSE)

write.table(DGY_,
            'C:/Gresham/tiny_projects/Project_Grace/supplemental_figures/Supplemental_Fig2C_Outliers_CNV.Only_DGY1740.txt', 
            sep='\t', row.names = FALSE, quote = FALSE)

points(DGY_hot$X1657, DGY_hot$X1740, col=rgb(1,0,0,0.5), pch = 0)

###
###
#pdf("C:/Gresham/tiny_projects/Project_Grace/supplemental_figures/Supplemental_Fig2C_all_combined_DGY1734.pdf")
combined <- merge(global_normalized_insertionPerGene, relative_depth_DNA_corrected_v3, by.x = "X", by.y = "Gene")
combined <- na.omit(combined)

cols <- rep(rgb(0,0,0,0.01),length(combined$X)) # initialize cols to black
cols[combined$DGY1734==2] <- rgb(1,0,0,0.5)
cols[combined$DGY1734==3] <- rgb(0,0,1,0.5)
cols[combined$DGY1734==4] <- rgb(0,1,1,0.5)

pchs <- rep(16, length(combined$X)) # initialize cols filled circle
pchs[combined$DGY1734==0] <- 3
pchs[combined$DGY1734==3] <- 17
pchs[combined$DGY1734==4] <- 18

ismax <- max(combined$X1657, combined$X1734)

# ismax <- max(combined$X1657, combined$X1728, combined$X1734, combined$X1736,
#              combined$X1740, combined$X1744, combined$X1747, combined$X1751,
#              na.rm =TRUE) 

plot(combined$X1657, combined$X1734, 
     col = cols, pch = pchs, xlim=c(0, ismax), ylim = c(0,ismax))

fit<-lm(combined$X1734~combined$X1657)
abline(fit)
summary(fit)

combined$standardized_residuals<-rstandard(fit)
res_cutoff<-2*sd(combined$standardized_residuals)+mean(combined$standardized_residuals)
DGY_hot<- subset(combined, abs(combined$standardized_residuals) > 2)

write.table(DGY_hot,
            'C:/Gresham/tiny_projects/Project_Grace/supplemental_figures/Supplemental_Fig2C_SigOutliers_DGY1734.txt', 
            sep='\t', row.names = FALSE, quote = FALSE)

write.table(combined,
            'C:/Gresham/tiny_projects/Project_Grace/supplemental_figures/Supplemental_Fig2C_Outliers_CNV_DGY1734.txt', 
            sep='\t', row.names = FALSE, quote = FALSE)


points(DGY_hot$X1657, DGY_hot$X1734, col=rgb(1,0,0,0.5), pch = 0)

DGY_ <- subset(combined, combined$DGY1734!=1)
fit<-lm((DGY_$X1734)~((DGY_$X1657)))
abline(fit, col='red')
summary(fit)

dev.off()

x<-subset(DGY_hot, DGY_hot$DGY1734!=1)
y<-subset(combined, combined$DGY1734!=1)
m<-subset(DGY_hot, DGY_hot$DGY1734==1)
n<-subset(combined, combined$DGY1734==1)

DGY_$standardized_residuals<-rstandard(fit)
DGY_hot<- subset(DGY_, abs(DGY_$standardized_residuals) > 2)

write.table(DGY_hot,
            'C:/Gresham/tiny_projects/Project_Grace/supplemental_figures/Supplemental_Fig2C_SigOutliers_CNV_DGY1734.txt', 
            sep='\t', row.names = FALSE, quote = FALSE)

write.table(DGY_,
            'C:/Gresham/tiny_projects/Project_Grace/supplemental_figures/Supplemental_Fig2C_Outliers_CNV.Only_DGY1734.txt', 
            sep='\t', row.names = FALSE, quote = FALSE)

###
###
###1736
#pdf("C:/Gresham/tiny_projects/Project_Grace/supplemental_figures/Supplemental_Fig2C_all_combined_DGY1736.pdf")
combined <- merge(global_normalized_insertionPerGene, relative_depth_DNA_corrected_v3, by.x = "X", by.y = "Gene")
combined <- na.omit(combined)

cols <- rep(rgb(0,0,0,0.01),length(combined$X)) # initialize cols to black
cols[combined$DGY1736==2] <- rgb(1,0,0,0.5)
cols[combined$DGY1736==3] <- rgb(0,0,1,0.5)
cols[combined$DGY1736==4] <- rgb(0,1,1,0.5)

pchs <- rep(16, length(combined$X)) # initialize cols filled circle
pchs[combined$DGY1736==0] <- 3
pchs[combined$DGY1736==3] <- 17
pchs[combined$DGY1736==4] <- 18

ismax <- max(combined$X1657, combined$X1736)

# ismax <- max(combined$X1657, combined$X1728, combined$X1734, combined$X1736,
#              combined$X1740, combined$X1744, combined$X1747, combined$X1751,
#              na.rm =TRUE) 

plot(combined$X1657, combined$X1736, 
     col = cols, pch = pchs, xlim=c(0, ismax), ylim = c(0,ismax))

fit<-lm(combined$X1736~combined$X1657)
abline(fit)
summary(fit)

combined$standardized_residuals<-rstandard(fit)
res_cutoff<-2*sd(combined$standardized_residuals)+mean(combined$standardized_residuals)
DGY_hot<- subset(combined, abs(combined$standardized_residuals) > 2)

write.table(DGY_hot,
            'C:/Gresham/tiny_projects/Project_Grace/supplemental_figures/Supplemental_Fig2C_SigOutliers_DGY1736.txt', 
            sep='\t', row.names = FALSE, quote = FALSE)

write.table(combined,
            'C:/Gresham/tiny_projects/Project_Grace/supplemental_figures/Supplemental_Fig2C_Outliers_CNV_DGY1736.txt', 
            sep='\t', row.names = FALSE, quote = FALSE)


points(DGY_hot$X1657, DGY_hot$X1736, col=rgb(1,0,0,0.5), pch = 0)

DGY_ <- subset(combined, combined$DGY1736!=1)
fit<-lm((DGY_$X1736)~((DGY_$X1657)))
abline(fit, col='red')
summary(fit)

dev.off()

x<-subset(DGY_hot, DGY_hot$DGY1736!=1)
y<-subset(combined, combined$DGY1736!=1)
m<-subset(DGY_hot, DGY_hot$DGY1736==1)
n<-subset(combined, combined$DGY1736==1)

DGY_$standardized_residuals<-rstandard(fit)
DGY_hot<- subset(DGY_, abs(DGY_$standardized_residuals) > 2)

write.table(DGY_hot,
            'C:/Gresham/tiny_projects/Project_Grace/supplemental_figures/Supplemental_Fig2C_SigOutliers_CNV_DGY1736.txt', 
            sep='\t', row.names = FALSE, quote = FALSE)

write.table(DGY_,
            'C:/Gresham/tiny_projects/Project_Grace/supplemental_figures/Supplemental_Fig2C_Outliers_CNV.Only_DGY1736.txt', 
            sep='\t', row.names = FALSE, quote = FALSE)

###1744
#pdf("C:/Gresham/tiny_projects/Project_Grace/supplemental_figures/Supplemental_Fig2C_all_combined_DGY1744.pdf")
combined <- merge(global_normalized_insertionPerGene, relative_depth_DNA_corrected_v3, by.x = "X", by.y = "Gene")
combined <- na.omit(combined)

cols <- rep(rgb(0,0,0,0.01),length(combined$X)) # initialize cols to black
cols[combined$DGY1744==2] <- rgb(1,0,0,0.5)
cols[combined$DGY1744==3] <- rgb(0,0,1,0.5)
cols[combined$DGY1744==4] <- rgb(0,1,1,0.5)

pchs <- rep(16, length(combined$X)) # initialize cols filled circle
pchs[combined$DGY1744==0] <- 3
pchs[combined$DGY1744==3] <- 17
pchs[combined$DGY1744==4] <- 18

ismax <- max(combined$X1657, combined$X1744)

# ismax <- max(combined$X1657, combined$X1728, combined$X1734, combined$X1736,
#              combined$X1740, combined$X1744, combined$X1747, combined$X1751,
#              na.rm =TRUE) 

plot(combined$X1657, combined$X1744, 
     col = cols, pch = pchs, xlim=c(0, ismax), ylim = c(0,ismax))

fit<-lm(combined$X1744~combined$X1657)
abline(fit)
summary(fit)

combined$standardized_residuals<-rstandard(fit)
res_cutoff<-2*sd(combined$standardized_residuals)+mean(combined$standardized_residuals)
DGY_hot<- subset(combined, abs(combined$standardized_residuals) > 2)

write.table(DGY_hot,
            'C:/Gresham/tiny_projects/Project_Grace/supplemental_figures/Supplemental_Fig2C_SigOutliers_DGY1744.txt', 
            sep='\t', row.names = FALSE, quote = FALSE)

write.table(combined,
            'C:/Gresham/tiny_projects/Project_Grace/supplemental_figures/Supplemental_Fig2C_Outliers_CNV_DGY1744.txt', 
            sep='\t', row.names = FALSE, quote = FALSE)


points(DGY_hot$X1657, DGY_hot$X1744, col=rgb(1,0,0,0.5), pch = 0)

DGY_ <- subset(combined, combined$DGY1744!=1)
fit<-lm((DGY_$X1744)~((DGY_$X1657)))
abline(fit, col='red')
summary(fit)

dev.off()

x<-subset(DGY_hot, DGY_hot$DGY1744!=1)
y<-subset(combined, combined$DGY1744!=1)
m<-subset(DGY_hot, DGY_hot$DGY1744==1)
n<-subset(combined, combined$DGY1744==1)

DGY_$standardized_residuals<-rstandard(fit)
DGY_hot<- subset(DGY_, abs(DGY_$standardized_residuals) > 2)

write.table(DGY_hot,
            'C:/Gresham/tiny_projects/Project_Grace/supplemental_figures/Supplemental_Fig2C_SigOutliers_CNV_DGY1744.txt', 
            sep='\t', row.names = FALSE, quote = FALSE)

write.table(DGY_,
            'C:/Gresham/tiny_projects/Project_Grace/supplemental_figures/Supplemental_Fig2C_Outliers_CNV.Only_DGY1744.txt', 
            sep='\t', row.names = FALSE, quote = FALSE)

###1747
#pdf("C:/Gresham/tiny_projects/Project_Grace/supplemental_figures/Supplemental_Fig2C_all_combined_DGY1747.pdf")
combined <- merge(global_normalized_insertionPerGene, relative_depth_DNA_corrected_v3, by.x = "X", by.y = "Gene")
combined <- na.omit(combined)

cols <- rep(rgb(0,0,0,0.01),length(combined$X)) # initialize cols to black
cols[combined$DGY1747==2] <- rgb(1,0,0,0.5)
cols[combined$DGY1747==3] <- rgb(0,0,1,0.5)
cols[combined$DGY1747==4] <- rgb(0,1,1,0.5)

pchs <- rep(16, length(combined$X)) # initialize cols filled circle
pchs[combined$DGY1747==0] <- 3
pchs[combined$DGY1747==3] <- 17
pchs[combined$DGY1747==4] <- 18

ismax <- max(combined$X1657, combined$X1747)

# ismax <- max(combined$X1657, combined$X1728, combined$X1734, combined$X1736,
#              combined$X1740, combined$X1744, combined$X1747, combined$X1751,
#              na.rm =TRUE) 

plot(combined$X1657, combined$X1747, 
     col = cols, pch = pchs, xlim=c(0, ismax), ylim = c(0,ismax))

fit<-lm(combined$X1747~combined$X1657)
abline(fit)
summary(fit)

combined$standardized_residuals<-rstandard(fit)
res_cutoff<-2*sd(combined$standardized_residuals)+mean(combined$standardized_residuals)
DGY_hot<- subset(combined, abs(combined$standardized_residuals) > 2)

write.table(DGY_hot,
            'C:/Gresham/tiny_projects/Project_Grace/supplemental_figures/Supplemental_Fig2C_SigOutliers_DGY1747.txt', 
            sep='\t', row.names = FALSE, quote = FALSE)

write.table(combined,
            'C:/Gresham/tiny_projects/Project_Grace/supplemental_figures/Supplemental_Fig2C_Outliers_CNV_DGY1747.txt', 
            sep='\t', row.names = FALSE, quote = FALSE)


points(DGY_hot$X1657, DGY_hot$X1747, col=rgb(1,0,0,0.5), pch = 0)

DGY_ <- subset(combined, combined$DGY1747!=1)
fit<-lm((DGY_$X1747)~((DGY_$X1657)))
abline(fit, col='red')
summary(fit)

dev.off()

x<-subset(DGY_hot, DGY_hot$DGY1747!=1)
y<-subset(combined, combined$DGY1747!=1)
m<-subset(DGY_hot, DGY_hot$DGY1747==1)
n<-subset(combined, combined$DGY1747==1)

DGY_$standardized_residuals<-rstandard(fit)
DGY_hot<- subset(DGY_, abs(DGY_$standardized_residuals) > 2)

write.table(DGY_hot,
            'C:/Gresham/tiny_projects/Project_Grace/supplemental_figures/Supplemental_Fig2C_SigOutliers_CNV_DGY1747.txt', 
            sep='\t', row.names = FALSE, quote = FALSE)

write.table(DGY_,
            'C:/Gresham/tiny_projects/Project_Grace/supplemental_figures/Supplemental_Fig2C_Outliers_CNV.Only_DGY1747.txt', 
            sep='\t', row.names = FALSE, quote = FALSE)

###1751
#pdf("C:/Gresham/tiny_projects/Project_Grace/supplemental_figures/Supplemental_Fig2C_all_combined_DGY1751.pdf")
combined <- merge(global_normalized_insertionPerGene, relative_depth_DNA_corrected_v3, by.x = "X", by.y = "Gene")
combined <- na.omit(combined)

cols <- rep(rgb(0,0,0,0.01),length(combined$X)) # initialize cols to black
cols[combined$DGY1751==2] <- rgb(1,0,0,0.5)
cols[combined$DGY1751==3] <- rgb(0,0,1,0.5)
cols[combined$DGY1751==4] <- rgb(0,1,1,0.5)

pchs <- rep(16, length(combined$X)) # initialize cols filled circle
pchs[combined$DGY1751==0] <- 3
pchs[combined$DGY1751==3] <- 17
pchs[combined$DGY1751==4] <- 18

ismax <- max(combined$X1657, combined$X1751)

# ismax <- max(combined$X1657, combined$X1728, combined$X1734, combined$X1736,
#              combined$X1740, combined$X1744, combined$X1747, combined$X1751,
#              na.rm =TRUE) 

plot(combined$X1657, combined$X1751, 
     col = cols, pch = pchs, xlim=c(0, ismax), ylim = c(0,ismax))

fit<-lm(combined$X1751~combined$X1657)
abline(fit)
summary(fit)

combined$standardized_residuals<-rstandard(fit)
res_cutoff<-2*sd(combined$standardized_residuals)+mean(combined$standardized_residuals)
DGY_hot<- subset(combined, abs(combined$standardized_residuals) > 2)

write.table(DGY_hot,
            'C:/Gresham/tiny_projects/Project_Grace/supplemental_figures/Supplemental_Fig2C_SigOutliers_DGY1751.txt', 
            sep='\t', row.names = FALSE, quote = FALSE)

write.table(combined,
            'C:/Gresham/tiny_projects/Project_Grace/supplemental_figures/Supplemental_Fig2C_Outliers_CNV_DGY1751.txt', 
            sep='\t', row.names = FALSE, quote = FALSE)


points(DGY_hot$X1657, DGY_hot$X1751, col=rgb(1,0,0,0.5), pch = 0)

DGY_ <- subset(combined, combined$DGY1751!=1)
fit<-lm((DGY_$X1751)~((DGY_$X1657)))
abline(fit, col='red')
summary(fit)

dev.off()

x<-subset(DGY_hot, DGY_hot$DGY1751!=1)
y<-subset(combined, combined$DGY1751!=1)
m<-subset(DGY_hot, DGY_hot$DGY1751==1)
n<-subset(combined, combined$DGY1751==1)

DGY_$standardized_residuals<-rstandard(fit)
DGY_hot<- subset(DGY_, abs(DGY_$standardized_residuals) > 2)

write.table(DGY_hot,
            'C:/Gresham/tiny_projects/Project_Grace/supplemental_figures/Supplemental_Fig2C_SigOutliers_CNV_DGY1751.txt', 
            sep='\t', row.names = FALSE, quote = FALSE)

write.table(DGY_,
            'C:/Gresham/tiny_projects/Project_Grace/supplemental_figures/Supplemental_Fig2C_Outliers_CNV.Only_DGY1751.txt', 
            sep='\t', row.names = FALSE, quote = FALSE)
```
